# Atoh1 is repurposed from neuronal to hair cell determinant by Gfi1 acting as a coactivator without redistributing Atoh1’s genomic binding sites

**DOI:** 10.1101/767574

**Authors:** Aida Costa, Lynn M. Powell, Abdenour Soufi, Sally Lowell, Andrew P. Jarman

**Affiliations:** Centre for Discovery Brain Sciences, Edinburgh Medical School, University of Edinburgh, Edinburgh EH8 9XD, UK; MRC Centre for Regenerative Medicine, Institute for Stem Cell Research, School of Biological Sciences, University of Edinburgh, Edinburgh EH16 4UU, UK; AgenTus Therapeutics, Chesterton, Cambridge, UK; Institute of Cell Biology, School of Biological Sciences, University of Edinburgh, Roger Land Building, Edinburgh EH9 3FF, UK

## Abstract

Although the lineage-determining ability of transcription factors is often modulated according to cellular context, the mechanisms by which such switching occurs are not well known. Using a transcriptional programming model, we found that Atoh1 is repurposed from a neuronal to an inner ear hair cell (HC) determinant by the combined activities of Gfi1 and Pou4f3. In this process, Atoh1 maintains its regulation of neuronal genes but gains ability to regulate HC genes. Pou4f3 enables Atoh1 access to genomic locations controlling the expression of sensory (including HC) genes, but Atoh1+Pou4f3 are not sufficient for HC differentiation. Gfi1 is key to the Atoh1-induced lineage switch, but surprisingly does not alter Atoh1’s binding profile. Gfi1 acts in two divergent ways. It represses the induction by Atoh1 of genes that antagonise HC differentiation, a function in keeping with its well-known repressor role in haematopoiesis. Remarkably, we find that Gfi1 also acts as a co-activator: it binds directly to Atoh1 at existing target genes to enhance its activity. These findings highlight the diversity of mechanisms by which one TF can redirect the activity of another to enable combinatorial control of cell identity.

## INTRODUCTION

During development, lineage commitment often relies on key transcription factors (TFs) that are necessary and sufficient to regulate the gene expression programme specific to a particular cell fate in a particular context. Such TFs, sometimes referred to as master regulators, include basic helix-loop-helix (bHLH) domain TFs such as MyoD, which determines muscle fate (Tapscott et al. 1988), and proneural factors, which determine neural fates (Masserdotti et al. 2016). Often, however, a TF can drive the specification of several seemingly unrelated cell types. How one TF can drive the formation of diverse cell types is a central question in developmental biology. Part of the answer is that master regulator TFs work in partnership with other TFs, with lineages defined by combinatorial codes of TFs. However, the mechanisms by which TFs modulate each other’s function are poorly understood. Recent ChIP-seq experiments have demonstrated that in at least some cases the same TF is targeted to different sites in the genome to determine alternative cellular identities according to composition of the combinatorial TF code. It remains an open question whether there are also cases where TF defines distinct lineages while remaining at the same genome locations.

It can be difficult to examine to what extent a particular TF is responsible for modifying DNA binding of another TF. Numerous cellular variables such as DNA accessibility, epigenetic landscape, TF cooperativity, cofactors, and enhancer activity influence the genome occupancy of TFs, making it difficult to unpick the relative influence of each of these variables when comparing two different lineage specification events. The advent of *in vitro* cell fate reprogramming has demonstrated that overexpression of specific TF combinations can be sufficient to drive specification of the same starting cells towards diverse cell types in the absence of other context-specific variables (Nakamori et al. 2017; Pfisterer et al. 2011; Zhao et al. 2015; Masserdotti et al. 2016). Therefore, cellular reprogramming provides a controlled system in which to investigate how TFs cooperate and modulate each other’s functions for cell fate specificity in the absence of other differences in cellular context (Aydin et al. 2019).

The bHLH-domain TF Atoh1 is an example of a cell lineage regulator that promotes the specification of diverse cell types in diverse tissue contexts (Jarman and Groves 2013). Atoh1-dependent cell types include: i) sensory hair cells (HCs) located in the inner ear (Bermingham et al. 1999; Chen et al. 2002; Pan et al. 2011) ii) a subset of neurons present in spinal cord and cerebellum (Klisch et al. 2011; Lai et al. 2011) iii) Merkel cells of the skin (Maricich et al. 2009), and iv) secretory cells in the intestine (Yang et al. 2001). Atoh1 function in HC formation is of particular interest because of its potential use in gene therapy to promote HC regeneration in sensorineural deafness. This arises from the finding that Atoh1 is not only necessary but also sufficient for HC formation in rodent models (Bermingham et al. 1999; Zheng and Gao 2000; Jarman and Groves 2013). However, *in vivo* HC regeneration resulting from experimentally supplied Atoh1 is very inefficient, but the factors limiting Atoh1’s function are poorly known (Costa et al. 2017).

There are several obstacles to understanding how Atoh1 specifically drives HC differentiation in the inner ear even though it has the ability to specify other cell types in other contexts. First, the spatial and temporal heterogeneity of embryonic tissues *in vivo* are difficult to dissect. Second, identifying *in vivo* binding sites by ChIP-seq requires large numbers of Atoh1-expressing cells. Atoh1 ChIP-seq analysis has been achieved in the context of cerebellar granule neurons (Lai et al. 2011) and intestinal crypt cells (Kim et al. 2014) but it is difficult in tissues where Atoh1-expressing cells are scarce such as the inner ear. For these reasons, the genome-wide binding occupancy profile of Atoh1 in HCs is yet to be determined.

Previously, we established the first direct programming strategy for generating a reliable supply of HC-like cells (induced HCs, iHCs) from mouse embryonic stem cells (mESCs) (Costa et al. 2015). Although forced expression of Atoh1 alone promoted neuronal differentiation, co-delivery of Gfi1 and Pou4f3 with Atoh1 promoted remarkably robust and efficient iHC differentiation. This suggests that Pou4f3 and Gfi1 can convert Atoh1 from a neuronal determinant to an HC determinant, in keeping with the fact that Pou4f3 and Gfi1 are both vital for HC survival and differentiation *in vivo* (Wallis et al. 2003a; Hertzano et al. 2004). However, it is not known how these two TFs modulate Atoh1’s function. Based on our current understanding, an attractive hypothesis would be that Pou4f3 and Gfi1 repress Atoh1’s neuronal transcriptional programme by simply redistributing Atoh1 binding away from neuronal genomic regulatory regions to new regions that control the expression of HC genes.

In this report, we used our TF programming strategy combined with genome-wide approaches (ChIP-seq and RNA-seq) to determine the effects of Gfi1 and Pou4f3 on Atoh1’s DNA binding and transcriptional activities. We demonstrate that Atoh1 can activate a neuronal programme in mESCs and confirm that Gfi1 and Pou4f3 convert Atoh1 to a sensory HC determinant. Surprisingly, however, Gfi1 and Pou4f3 do not repress Atoh1’s neuronal programme during this process, but instead confer on Atoh1 an additional capacity for HC gene regulation. Pou4f3 and Gfi1 achieve this in different ways. Pou4f3 recruits Atoh1 to new genomic sites, but by itself this results in the activation of a generic sensory program and reinforcement of neuronal differentiation. Unlocking the HC programme requires Gfi1, but it does not act by altering the DNA binding profiles of Atoh1 or Pou4f3. Instead, Gfi1 has two functions. First, it represses inappropriate gene expression, consistent with previously described roles in hematopoiesis (van der Meer et al. 2010). Second, Gfi1 interacts with Atoh1 at existing sites as a potent transcriptional co-activator to augment both neuronal and HC-specific sensory programs. Overall, these findings demonstrate how repurposing lineage determining TFs is not necessarily achieved through major change in their binding profiles but can be accomplished in part through more subtle effects such as transcriptional amplification of pre-existing binding sites. Therapeutically, these results suggest new avenues to overcome Atoh1’s limitations in promoting HC regeneration.

## RESULTS

### Gfi1 and Pou4f3 facilitate HC gene activation without repressing Atoh1’s neuronal programme

To investigate how Gfi1 and Pou4f3 modulate the lineage determining activity of Atoh1, we engineered mouse embryonic stem cell (ESC) lines that allow for dox-inducible co-expression of Gfi1, Pou4f3 and Atoh1 (iGPA) or Atoh1 alone (iAtoh1). In each polycistronic cassette, the transgenes were separated by 2AP and end with an mVenus fluorescent reporter (Fig 1A). To facilitate immunoprecipitation for subsequent ChIP-seq analysis, FLAG and MYC tags were appended to the C-terminus of Atoh1 and N-terminus of Pou4f3 respectively (Fig. 1A). This double-tagged iGPA line was able to induce iHC differentiation with similar efficiency to that of the previously described untagged iGPA counterpart (iGPA-Myo7a:mVenus line (Costa et al. 2015) (Supplemental Fig. S1). We also attempted to tag Gfi1 with an HA epitope, resulting in a triple-tagged iGPA cell line, but this line did not support iHC differentiation upon transgene induction, and therefore the double-tagged iGPA line was used in subsequent experiments (Supplemental Fig. S1A).

**Figure 1.**
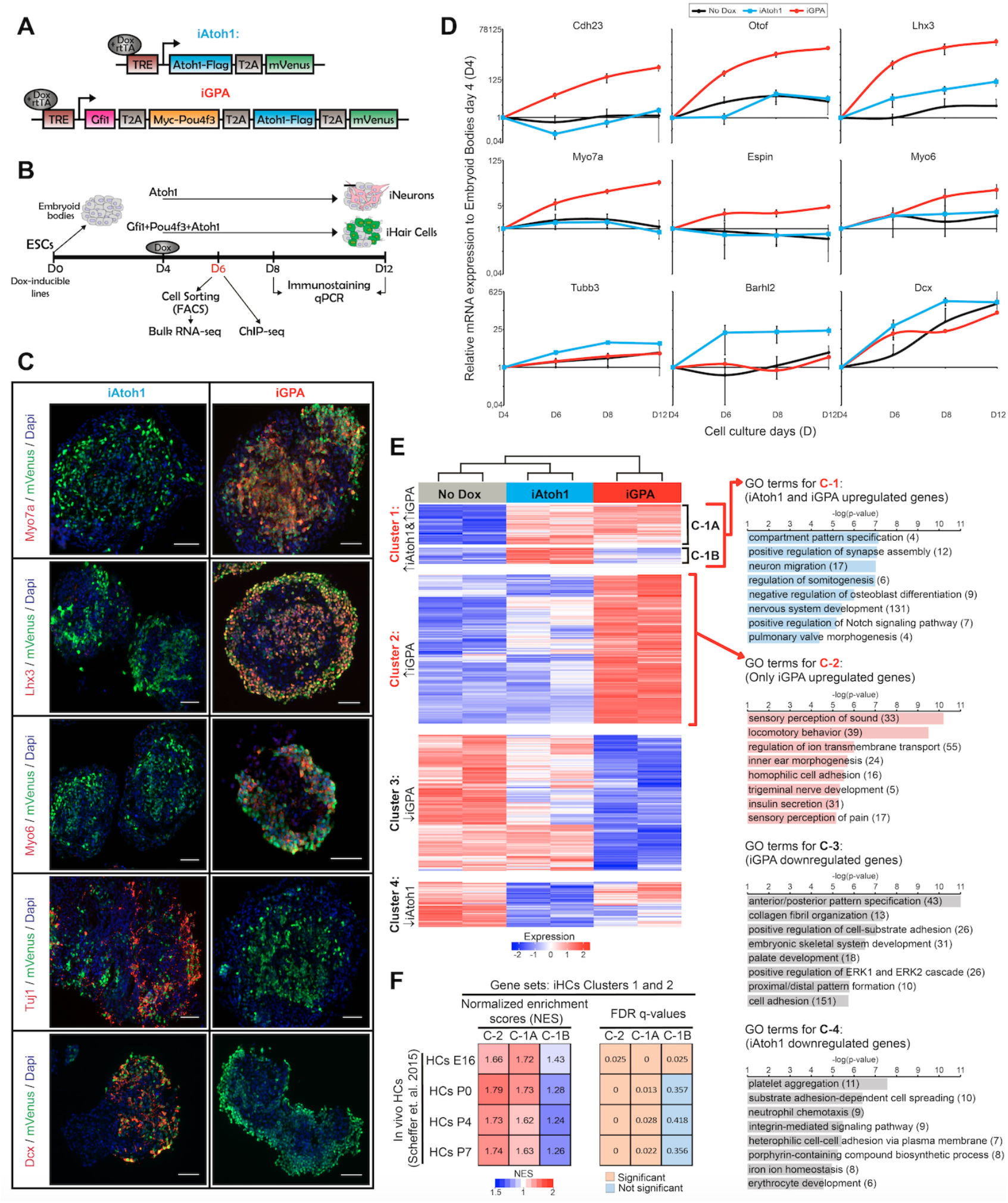
Atoh1 drives neuronal differentiation while Gfi1+Pou4f3+Atoh1 drives HC differentiation. (A) Dox-inducible tagged TF constructs used. (B) Schematic protocol for TF induction in EBs. (C) Expression of HC (Myo7a, Lhx3, Myo6) and neuronal (Tuj1, Dcx) markers (red) in induced EBs relative to mVenus (green) and DAPI (blue). (D) qPCR in EBs for HC (Cdh23, Otof, Lhx3, Myo7a, Espin, Myo6) and neuronal (Tubb3, Barhl2, Dcx) markers. (E) Cluster analysis and enriched GO terms for transcriptomes of iAtoh1 and iGPA EBs relative to uninduced EBs. Clusters 1-4 are shown; Cluster 1 is divided into 1A and 1B. (F) Normalised enrichment scores for clusters 1A, 1B and 2.

To induce differentiation, embryoid bodies (EBs) were generated from iGPA and iAtoh1 lines and were then treated with Dox for 2, 4 and 8 days (Fig. 1B). Gene expression analysis of mVenus+ cells revealed that while iGPA lines induced the expression of HC markers, iAtoh1 cells predominantly activated general neuronal genes (Fig. 1C,D). This confirms that forced expression of Atoh1 alone promotes neuronal differentiation in agreement with previous studies using *in vitro* ESC differentiation assays (Ebeid et al. 2017; Sagal et al. 2014; Srivastava et al. 2013).

We set out to test the initial hypothesis that Gfi1 and/or Pou4f3 repress Atoh1’s neuronal programme during iHC differentiation. We performed RNA-seq in FACS-purified mVenus^+^ cells harvested 48 h after Dox induction and identified genes differentially expressed (DE) compared to non-dox treated EBs. As expected, DE genes in iGPA cells display an HC signature, while those in iAtoh1 cells showed a neuronal signature (Supplemental Fig. S2A,B).

Unsupervised clustering of all DE genes (2818 genes) revealed four prominent clusters (Fig. 1E). Cluster 1 and 2 contained genes that are enriched in Dox-treated EBs relative to non-Dox-treated controls and these are mainly associated with neuronal and HC identity. On the other hand, genes within cluster 3 and 4 were mainly depleted relative to non-Dox controls and are not specifically associated with neurons or HCs. We focused our subsequent analysis on genes within cluster 1 and 2. Cluster 2 contains genes that were exclusively enriched in iGPA cells and are mainly associated with HC development, ion transport and sensory neuronal development (Fig. 1E). Cluster 1 contains genes associated with neuronal development, which as expected were enriched in iAtoh1 cells (Fig. 1E). Surprisingly, most of these genes (cluster 1A: 363 out of 511 genes) were also active in iGPA cells despite these cells not displaying an apparent neuronal identity. Only a small subset of neuronal genes including Barh12, Tubb3, and Dcx were specifically activated in iAtoh1 cells and not in iGPA cells (cluster 1B; Supplemental Fig. S2C). These findings indicate that iGPA cells differentiate to the HC lineage while retaining a large part of the neuronal program that is induced in iAtoh1 cells.

To determine whether this Atoh1 neuronal programme is also active during HC differentiation *in vivo*, we performed gene set enrichment analyses of clusters 1A, 1B and 2 on RNA-seq data previously obtained from HCs of the mouse inner ear (Scheffer et al. 2015). This revealed greater and more significant enrichments of clusters 1A and 2 genes compared to cluster 1B genes in both embryonic and postnatal HC transcriptomes (Fig. 1F). The fact that genes in clusters 1A and 2 are equally enriched in native HCs at early stages of differentiation (HCs E16, Fig. 1F) suggests that the dual activation of both neuronal and sensory differentiation programs is a feature of HC development *in vivo*. Conversely, the subset of Atoh1 neuronal targets that are repressed in *in vitro* iHCs (cluster 1B) are also less abundant during early HC differentiation stages *in vivo*. Taken together, these data indicate that genes activated in response to Atoh1 include one set activated in both neuronal and HC contexts (cluster 1A) and another set activated in only the HC context (cluster 2). In addition, a smaller subset of genes activated by Atoh1 are repressed in the HC context (cluster 1B).

We conclude that, contrary to our initial hypothesis, the modulation of Atoh1 by Gfi1/Pou4f3 entails activation of a new HC gene expression programme but without concomitant widespread repression of Atoh1’s neuronal programme.

### Gfi1 is critical for the switch from neuronal to HC fate

To investigate the individual contributions of Pou4f3 and Gfi1 to modulating Atoh1-induced differentiation, we generated cell lines that allow inducible expression of every combination of the three TFs (Fig. 2A). We compared global transcriptional changes using hierarchical clustering and principle component analysis. On their own, Gfi1 or Pou4f3 possess little capacity to modify gene expression (iGfi1 and iPou4f3 in Fig. 2B,C). Surprisingly the other TF combinations resulted in gene expression programmes that clustered into two distinct groups correlating with the presence or absence of Gfi1 (Fig. 2B,C).

**Figure 2.**
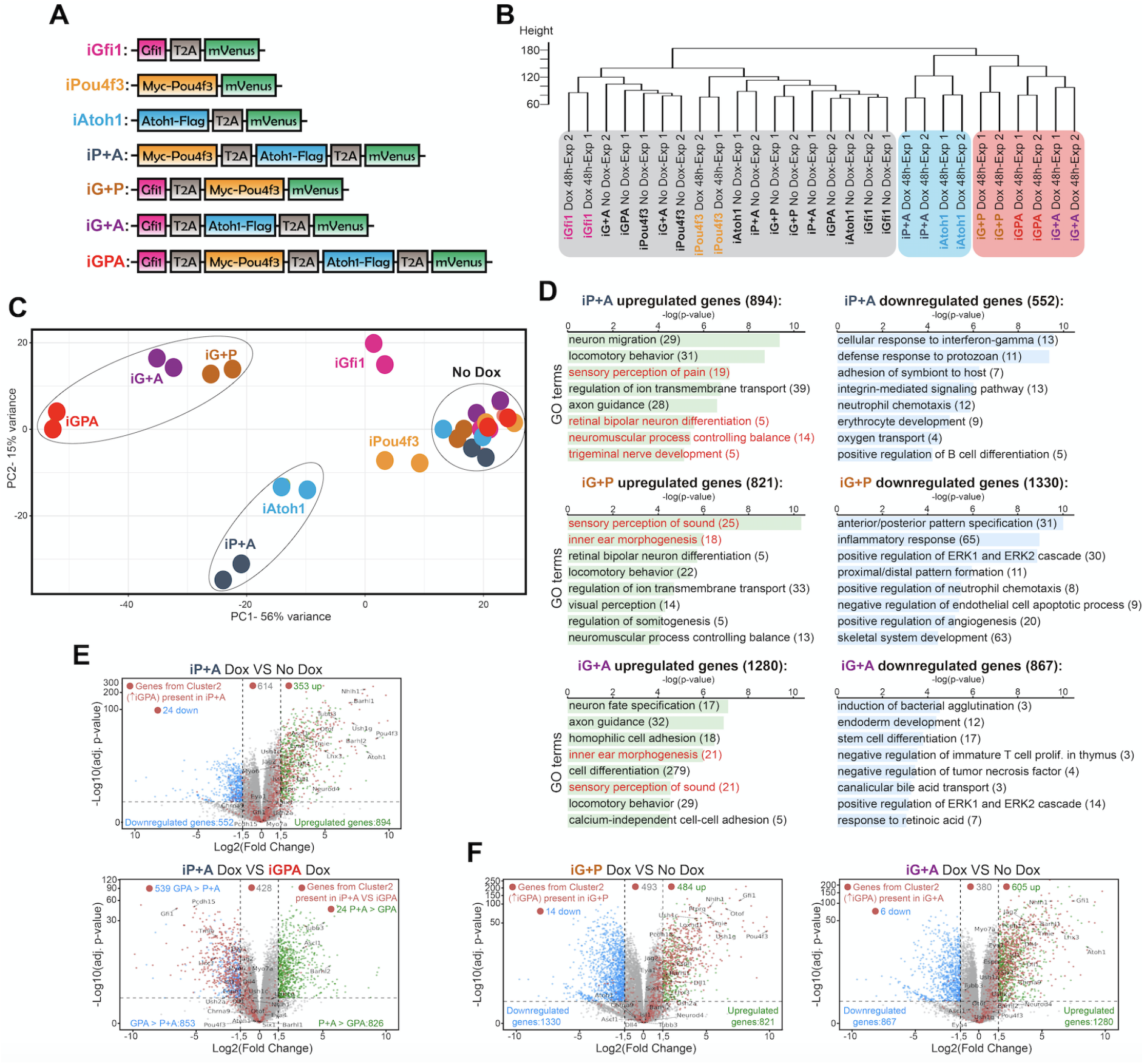
Transcriptome analysis of differentiation supported by TF combinations reveals Gfi1 is critical for HC switch. (A) Dox-inducible tagged TF constructs used. (B) Cluster analysis of transcriptomes from EBs induced to express different TF combinations. (C) Principal component analysis of transcriptomes. (D) Enriched GO terms in transcriptomes. (E) Volcano plot of transcriptome expression levels for iP+A. (F) Volcano plot of transcriptome expression levels for iG+P and iG+A.

The transcriptome of iP+A cells resembles that of iAtoh1 cells rather than that of iGPA cells, suggesting a shared neuronal identity (Fig. 2B and C). Consistent with this, Atoh1 + Pou4f3 were unable to induce HC markers in EBs even after 8 days of Dox treatment. Rather, general neuronal markers were upregulated in iP+A cells even more strongly than in iAtoh1 cells (Supplemental Fig. S3A). Therefore, Pou4f3 alone cannot convert Atoh1 from a neuronal determinant to a HC determinant.

Pou4f3 alone is, however, able to modify Atoh1’s neuronal programme. Most strikingly, the genes enriched in iP+A but not iAtoh1 are involved in the development of sensory neurons (Fig. 2D) and are associated with sensory functions such as ‘perception of pain’ (Supplemental Fig. S3B). Although iP+A did not induce HC differentiation, we were able to detect 35% of genes from the HC-related DE gene cluster 2 amongst the genes upregulated by iP+A (Fig. 2E). However, their expression remained considerably lower than in iGPA cells (Fig. 2E) and seemed to be insufficient to trigger the activation of the HC differentiation program. Altogether, these data suggest that the co-expression of Atoh1 and Pou4f3 promotes the activation of a sensory neuronal program, not a functional HC programme.

In contrast, expression profiles of iG+P and iG+A cells cluster strongly with iGPA cells, suggesting that Gfi1 in combination with either Atoh1 or Pou4f3 or both drives cells to a common differentiation path related to HCs (Fig. 2B,C). The presence of Gfi1 therefore contributes to a major shift in gene expression when it is combined with Atoh1 and/or Pou4f3 (Fig. 2C). This differentiation pathway is characterised by activation of genes required for sensory perception of sound, consistent with HC identity (Fig. 2D,F). These observations support a model in which Gfi1 dramatically modulates the transcriptional activity of both Atoh1 and Pou4f3 to enable HC differentiation.

The ability of Gfi1+Pou4f3 to drive expression of HC genes in the absence of exogenous Atoh1 was a surprise. We therefore examined the efficiency of HC differentiation in iG+P cells using immunostaining and qPCR for HC and neuronal markers. This confirmed that Gfi1 abolishes the neurogenic activities of both Atoh1 and Pou4f3 and enables each of them instead to initiate an HC differentiation programme (Fig. 3A–C). However, we found that HC differentiation was not as efficient in either iG+P or iG+A cells as it was in iGPA cells (Fig. 1C). For example, the HC marker Lhx3 was confined to a minor subset of mVenus+ cells in both iG+P and iG+A lines (Fig. 3A). Furthermore, iG+P and iG+A cells were unable to develop Espin+ hair bundle-like structures, in contrast to iGPA cells which display a clear polarised Espin signal (Fig. 3D–F; Supplemental Fig. S1A). Instead, Espin staining was spread throughout the cytoplasm of the mVenus+ iG+P and iG+A cells. This suggests that despite the expression of hair bundle genes (Fig. 3C), key components necessary for organization of these hair-bundle-like protrusions are likely to missing from iG+P and iG+A cells.

**Figure 3.**
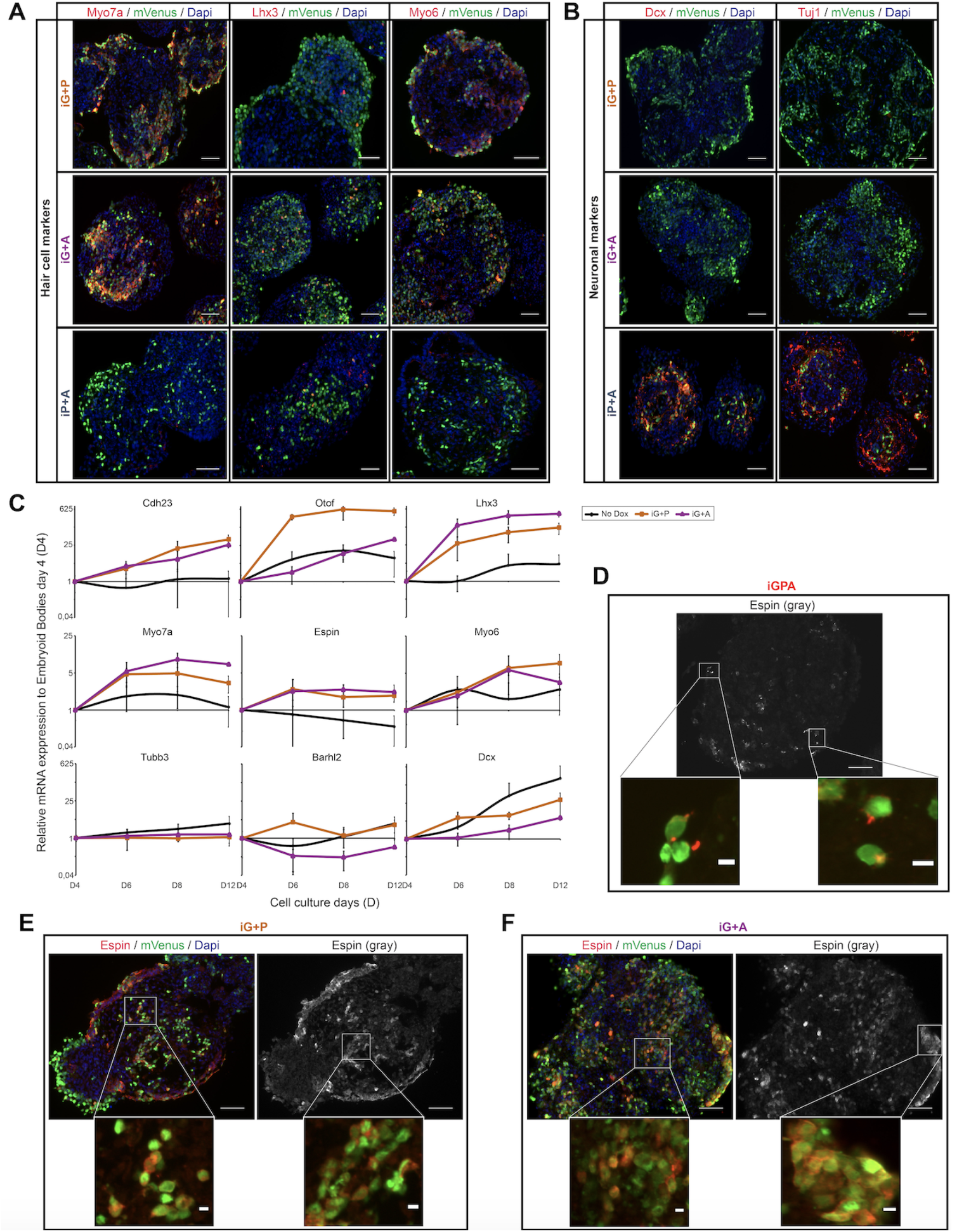
Gfi1 with Atoh1 or Pouf43 promotes HC gene expression but cellular differentiation requires all three TFs. (A) Expression of HC markers (Myo7a, Lhx3, Myo6, red) in induced EBs relative to mVenus (green) and DAPI (blue). (B) Expression of neuronal markers (Dcx, Tuj1, red) in induced EBs. (C) qPCR in EBs for HC (Cdh23, Otof, Lhx3, Myo7a, Espin, Myo6) and neuronal (Tubb3, Barhl2, Dcx) markers. (D) Espin expression in polarised stereocilia-like outgrowths in iGPA cells. (E,F) Espin expression is not polarised in iG+P (E) or iG+A (F) EBs.

In summary, we find that complete activation of the iHC differentiation programme requires all three TFs. Gfi1 plays a critical role by imposing a dramatic change on the transcriptional activity of both Atoh1 and Pou4f3, thereby allowing these TFs to activate a partial HC differentiation program individually, or a more complete HC programme together.

### Atoh1 directly regulates both neuronal and sensory genes

Having identified genes upregulated in response to Atoh1 in the contexts of neuronal or HC differentiation, we asked which of these genes might be directly regulated by Atoh1. Taking advantage of the FLAG-tagged Atoh1, we carried out Atoh1 ChIP-Seq on both iAtoh1 and iGPA lines after inducing transgene expression in EBs for 48 h. 15568 genomic sites were enriched for Atoh1 in iAtoh1 cells and 262231 sites in iGPA cells. Together, these sites could be divided into three groups (Supplemental Fig. S4A): iAtoh1-unique (7279 sites), iAtoh1/iGPA-common (8056 sites), and iGPA-unique (17851 sites) (Fig. 4A). We examined whether these differences in Atoh1 binding explain how Atoh1 switches from neuronal to HC determinant.

**Figure 4.**
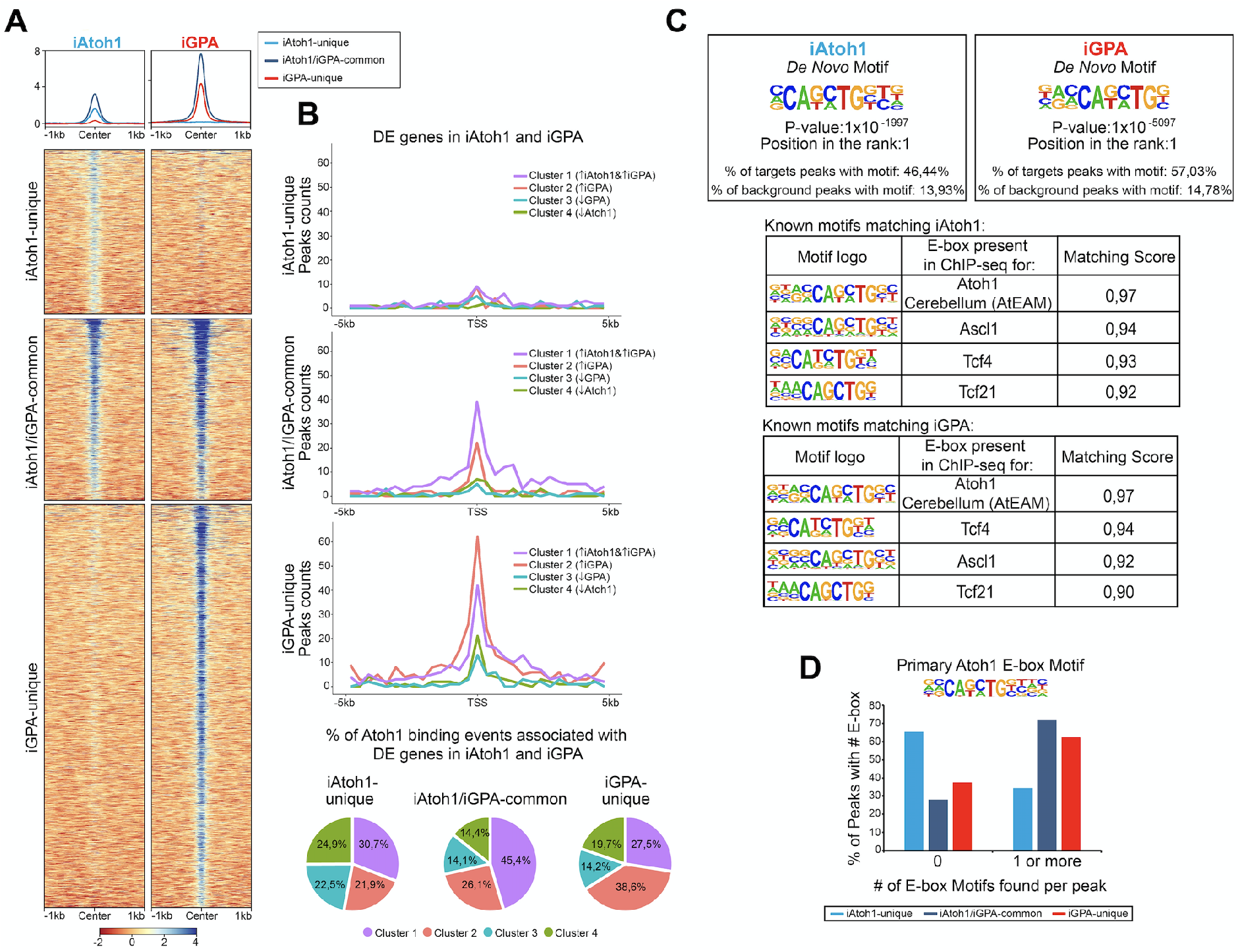
Gfi1+Pou4f3 promote Atoh1 binding to its preferred motif at new genomic locations without inhibiting binding to its original targets. (A) Heat maps of three classes of Atoh1 ChIP-seq peak in iAtoh1 and iGPA EBs. (B) Atoh1 binding peaks associated with DE clusters in iAtoh1 and iGPA EBs. (C) Sequence motifs enriched in Atoh1 binding peaks from iAtoh1 and iGPA EBs. Upper: highest *de novo* motif detect is an E box (CANNTG); lower: best scoring known E box motif is the Atoh1 binding motif previously detected in cerebellum (AtEAM). (D) Number of AtEAM motifs in binding peaks associated with the three classes of Atoh1 binding site in (A).

To determine whether these groups are associated with the DE gene clusters identified above, we integrated Atoh1 genomic location analyses with these DE gene clusters by annotating the peaks to their closest TSS. First, the iAtoh1-unique group of sites showed relatively low binding enrichments and were not preferentially associated with any particular DE gene cluster (Fig. 4A,B). We therefore surmise that this group likely reflects weak and non-functional binding of Atoh1. We next examined iAtoh1/iGPA-common binding sites. These were predominantly associated with DE cluster 1 (enriched in neuronal genes), indicating that Atoh1 directly binds and activates neuronal genes regardless of the presence or absence of Pou4f3 and Gfi1. Consistent with this, iAtoh1/iGPA-common sites included enhancers that were previously shown to be enriched for Atoh1 in the developing cerebellum (Klisch et al. 2011) intestinal crypts (Kim et al. 2014) and spinal cord (Lai et al. 2011) (Supplemental Fig. S4B), validating our ChIP-seq strategy using the FLAG tag. Finally, the iGPA-unique sites were predominantly associated with genes of DE cluster 2 (enriched in HC genes) (Fig. 4B), suggesting that Atoh1 directly binds and activates HC genes but only in the presence of Pou4f3/Gfi1, although is not possible to determine from these data whether it is Pou4f3 or Gfi1 or both that enable Atoh1 to bind HC genes.

In *de novo* motif analysis of Atoh1 binding peaks, the best hit was an E box motif that matched most closely the AtEAM variant previously identified from Atoh1 binding sites in the cerebellum^6^ (Fig. 4C; Supplemental Fig. S5). More than 60% of iAtoh1/iGPA-common and iGPA-unique peaks contained one or more AtEAM E-box motifs compared to just 34.5% of iAtoh1-unique peaks (Fig. 4D). This is consistent with the previous conclusion that the majority (65.5%) of iAtoh1 unique peaks largely represent low affinity and non-specific interactions. Altogether, these data suggest that Pou4f3 and/or Gfi1 do not interfere with Atoh1 binding at neuronal targets with its preferred E-box motif, but enable Atoh1 to bind to HC targets containing its preferred E-box motif, while depleting Atoh1 from weakly-bound sites which for the most part lack the E-box motif.

### Atoh1 recruitment to new targets is largely driven by Pou4f3, not Gfi1

We next determined the relative importance of Pou4f3 and Gfi1 in directing Atoh1 to new binding sites during iHC differentiation. We performed ChIP-seq for Atoh1, and also for Pou4f3, after induction of all combinations of the three TFs. PCA and hierarchical clustering revealed that global Atoh1 binding is distinctly altered by the presence of Pou4f3 (iP+A and iGPA binding cluster distinctly from iAtoh1 binding) (Fig. 5A,B). In contrast, the presence of Gfi1 does not strongly affect the profile of Atoh1 binding, regardless of the presence or absence of Pou4f3 (Atoh1 profiles in iAtoh1 vs iG+A and in iGPA vs iP+A, Fig. 5A,B). Therefore, despite Gfi1’s importance for promoting the HC differentiation programme, the presence of Gfi1 did not make a large difference to the binding pattern of Atoh1.

**Figure 5.**
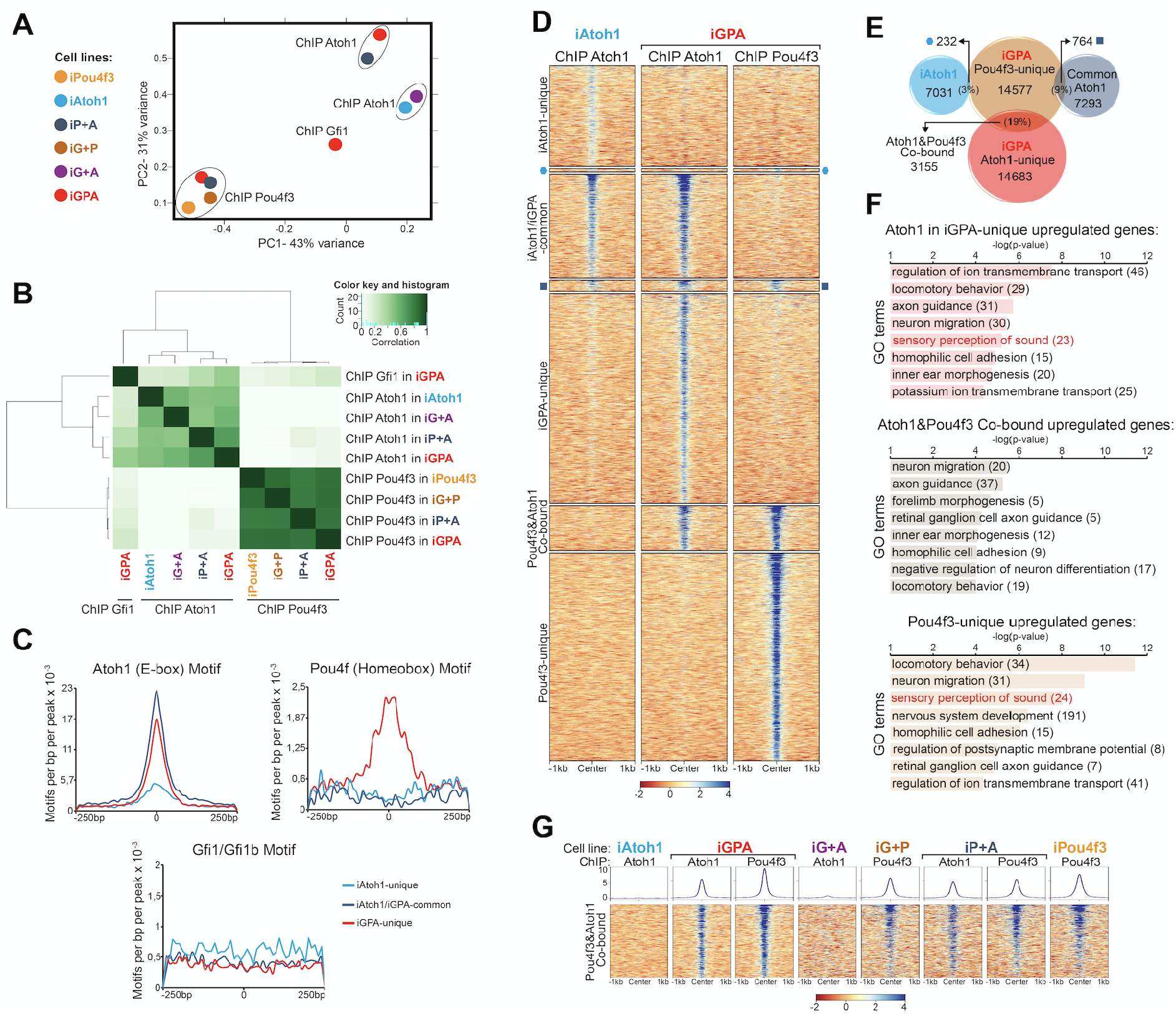
Pou4f3 recruits Atoh1 to new binding locations but not vice versa. (A) Principal component analysis of ChIP-seq binding peaks for Atoh1, Pou4f3 and Gfi1 in EBs expressing combinations of the three TFs (key on LHS). (B) Cluster analysis of ChIP-seq binding peaks. (C) Atoh1, Pou4f and Gfi1 DNA motifs associated with Atoh1 binding peaks, separated according to the three classes of Atoh1 site (Fig. 4A). (D) Heat maps of ChIP-seq enrichment for Atoh1 and Pou4f3 binding. (E) Venn diagram of Pou4f3 binding peaks in iGPA EBs (orange) relative to Atoh1 binding peaks separated according to the three classes in Fig. 4A. (F) Enriched GO terms associated with DE genes linked to Atoh1 and/or Pou4f3 binding peaks. (G) Heat maps of Atoh1 and Pou4f3 ChIP-seq enrichment at Atoh1/Pou4f3 cobound locations in different induced EBs.

In contrast to Atoh1, Pou4f3 binding profiles in all conditions cluster tightly together, suggesting that Pou4f3 binding is not strongly affected by the presence of either Atoh1 or Gfi1 (Fig. 5A,B). Taken together our data suggest that the binding of Atoh1 to new sites during HC differentiation depends directly or indirectly on Pou4f3 function to a large extent. In contrast, and contrary to expectation, Gfi1 allows Atoh1 and Pou4f3 to activate a new transcriptional programme largely without redirecting their binding to new loci.

### Pou4f3 directly recruits Atoh1 to a subset of new neuronal targets

The genomic binding profile of Pou4f3 is quite different from Atoh1 (Fig. 5A,B), suggesting that much of the effect of Pou4f3 on Atoh1 binding redistribution might be indirect. To investigate the possibility of direct interactions, we searched for the enrichment of motifs in addition to E boxes in each of the three different groups of Atoh1 peaks. A homeobox motif bound by Pou4f TFs was the most significant secondary motif found in iGPA-unique peaks but was absent from the other peak categories (Fig. 5C and Supplemental Fig. S5B). This is consistent with the possibility that Pou4f3 might directly promote Atoh1 binding to new regulatory elements. To explore this, we performed clustering of the genomic locations for Atoh1 and Pou4f3 in the different TF combinations (Fig. 5D,E). This reveals that Atoh1 co-binds with Pou4f3 at a subset of sites (3155 sites in cluster_3). The majority of these iGPA co-bound sites contained both Pou4f and E-boxes motifs with similarly high abundance (Supplemental Fig. S6A,B), suggesting that co-binding of Atoh1 and Pou4f3 is dependent on the presence of both their specific motifs in the same regulatory element.

Notably, these co-bound sites are bound by Pou4f3 in all conditions (and therefore are not dependent on the presence of Atoh1), whereas binding of Atoh1 is prominent in iGPA and iP+A but not iAtoh1 or iG+A cells (Fig. 5G). We conclude that Atoh1 binding is dependent on Pou4f3 but not vice versa, consistent with the idea that Pou4f3 recruits Atoh1 to these new sites.

To explore whether the cluster of Atoh1+Pou4f3 co-bound sites includes genes activated upon HC differentiation, we integrated the RNA-seq and ChIP-seq datasets. We annotated the peaks in each category and performed GO analyses only for genes previously identified as DE in iGPA cells. This analysis revealed that DE genes with GO terms related to HC differentiation were mostly bound independently by Atoh1 or Pou4f3 (unique sites) with only a minor fraction of HC genes co-bound by both TFs (Fig. 5F). Instead GO terms related to neuronal migration and axon guidance were most significant among the co-bound sites (Fig. 5F). Overall, these data suggest that Pou4f3 directly recruits Atoh1 to new loci, but these are associated with neuronal rather than HC differentiation. In contrast, recruitment of Atoh1 to HC loci by Pou4f3 therefore appears to be largely indirect.

### Gfi1 co-binds with Atoh1 at a subset of sites

Having shown that Gfi1 is critical for converting Atoh+Pou4f3 from neuronal determinants to HC determinants, it was surprising that it does not appear to redistribute Atoh1 or Pou4f3 binding. Moreover, Atoh1 binding peaks in iGPA cells are not associated with Gfi1 motifs (Fig. 5C). We therefore considered a model in which Gfi1 functions in parallel to and independently of Atoh1/Pou4f3. One prediction of this model is that Gfi1 does not co-bind with Atoh1 or Pou4f3 during HC differentiation. To test this, we performed ChIP-seq for Gfi1 in iHCs generated from the iGPA line. Since we were unable to epitope tag Gfi1, we relied instead on a Gfi1 antibody for immunoprecipitation. This antibody gave a relatively low binding enrichment over the input DNA, resulting in only 3003 detectable peaks. Nevertheless, Gfi1-unique peaks (those that did not overlap with Atoh1 or Pou4f3) were strongly enriched in Gfi1 DNA motifs (see below), pointing to the specificity of binding.

We analysed these Gfi1 binding peaks for overlap with the other two TFs. To our surprise, 43% of these peaks overlapped with those of Atoh1 Fig. 6A. Furthermore, Pou4f3 peaks overlapped with Gfi1 peaks mostly only where Atoh1 peaks were also present. Motif analysis revealed that Atoh1-Gfi1 co-bound peaks are not enriched in Gfi1 DNA motifs (Fig. 6B,C). This corroborates our previous finding that Atoh1 and Pou4f3 peaks lack Gfi1 binding motifs (Supplemental Fig. S6A). Instead, E-boxes featured prominently in those Gfi1 peaks that coincide with Atoh1 peaks. Pou4f motifs were also detectable, but only in peaks in common to all three TFs (Fig. 6B,C). Together, these data suggest that Gfi1 is recruited to a set of loci through interaction with Atoh1 rather than through specific DNA binding. In support of this hypothesis we were able to recover Myc-tagged Atoh1 from an *in vitro* pulldown assay using a recombinant GST-Gfi1 protein (Fig. 6G).

**Figure 6.**
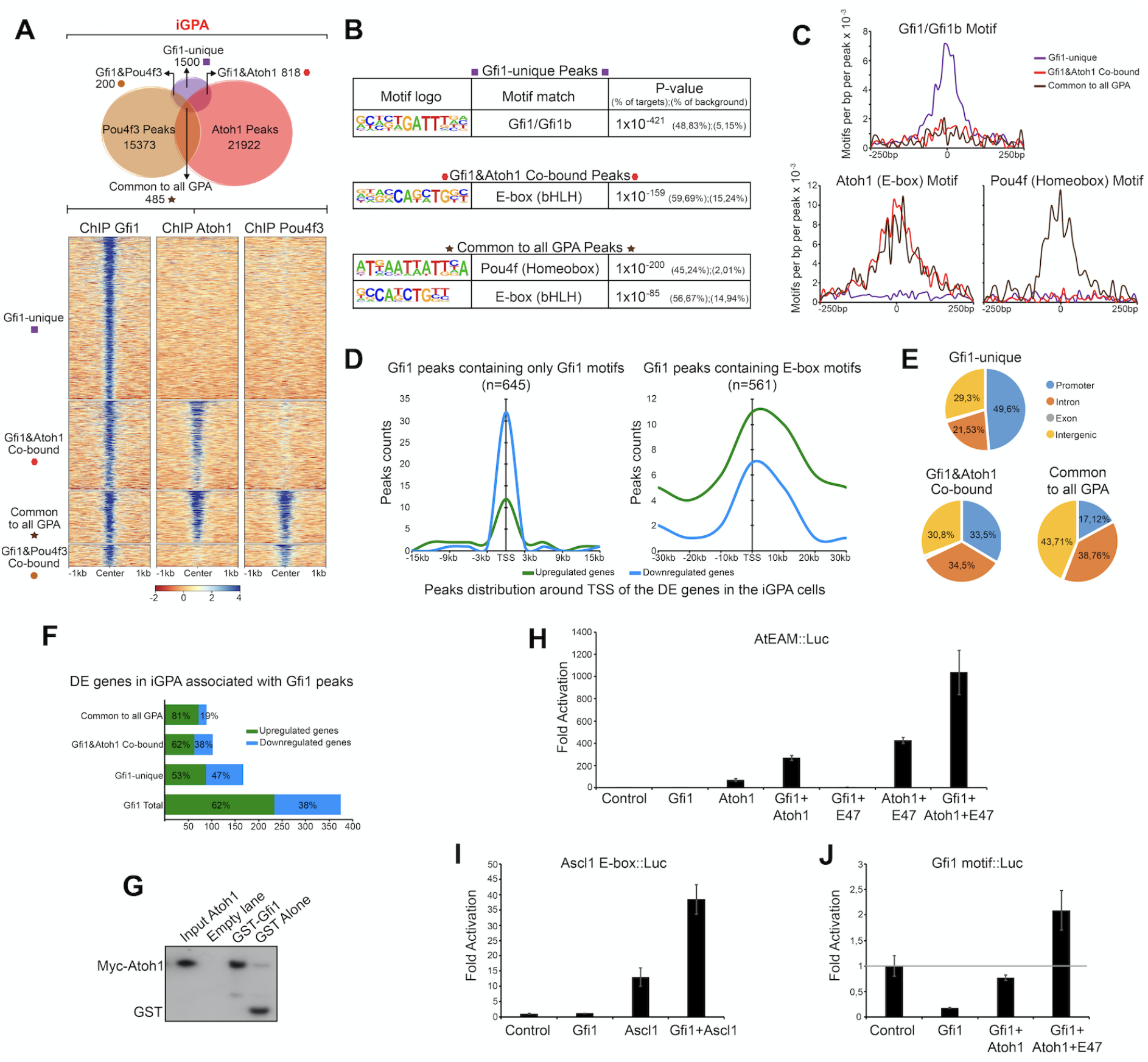
Gfi1 unique sites are enriched in Gfi1 motifs but Gfi1/Atoh1 cobound sites are enriched in E boxes. (A) Upper: Venn diagram of Gfi1, Pou4f3 and Atoh1 binding locations in iGPA cells. Lower: Heat maps of Gfi1, Atoh1 and Pou4f3 ChIP-seq enrichment at locations in iGPA EBs. (B) Motifs associated with Gfi1 binding peaks in iGPA EBs, separated according to (A). (C) Locations of motifs at the Gfi1 binding peaks in iGPA cells, separated according to (A). (D) Gfi1 binding peaks distributed around the TSS of DE genes in iGPA EBs. (E) Pie charts of Gfi1 binding peak location in associated genes. (F) Numbers of DE genes associated with Gfi1 binding peaks in iGPA cells. (G) GST pull-down analysis of Myc-tagged Atoh1 with GST-tagged Gfi1. Western blot probed with anti-Myc antibody. (H) Expression analysis of luciferase reporter gene under control of multimerised Atoh1 (AtEAM) motifs in cells cotransfected with expression vectors for Gfi1, Atoh1 and/or E47 (control is cotransfection with empty expression vector). Fold activation relative to control. The data with E47 are from a separate experiment from the data without E47. (I) Similar expression analysis of luciferase reporter gene under control of multimerised Ascl1 motifs. (J) Similar expression analysis of luciferase reporter gene under control of multimerised Gfi1 binding motifs.

In addition to the overlapping Gfi1 peaks above, 50% of Gfi1 binding sites did not overlap with those of Atoh1 or Pou4f3. In contrast to the overlapping peaks, these ‘Gfi1-unique’ sites were enriched in Gfi1 motifs (Fig. 6B,C). Together, these findings raise the possibility that Gfi1 binds to one set of target sites directly and independently but is recruited to another set of loci through interaction with Atoh1.

### Gfi1 regulates HC differentiation through both repression and Atoh1 co-activation

Gfi1 protein contains a SNAG domain which recruits co-repressors (Velinder et al. 2016; Zweidler-Mckay et al. 1996; Grimes et al. 1996), and it acts as a major transcriptional repressor during haematopoiesis (Möröy et al. 2015; Möröy and Khandanpour 2011; van der Meer et al. 2010). However, the *Drosophila* orthologue of Gfi1 (Senseless, SENS) (Nolo et al. 2000) has been reported to have dual functionality: SENS can act as a DNA-binding transcriptional repressor but also as a transcriptional co-activator of proneural TFs (including Atonal), enhancing their ability to stimulate sensory gene expression (Acar et al. 2006; Jafar-Nejad et al. 2003a; Powell et al. 2008; 2012). Our observation of two distinct populations of Gfi1 sites in iHCs suggests that Gfi1 may similarly have dual functionality. We hypothesised that i) Gfi1 represses genes that it binds to through its own motif independently of Atoh1, and ii) interaction between Gfi1 and Atoh1 at Atoh1-dependent sites switches Gfi1 from a repressor to a co-activator.

To test this hypothesis, we separated the Gfi1 peaks into two groups: one containing only Gfi1 motifs and the other containing E-box motifs, and assessed whether these were associated with genes that were activated or with genes that are repressed during iHC differentiation of iGPA cells. Consistent with Gfi1’s known role in hematopoiesis, peaks with Gfi1 motifs appeared to be strongly associated with gene repression, albeit only when located near to the TSS of the assigned genes (Fig. 6D). In support of this, we observed that 49.6% of Gfi1-unique sites are located near promoters and around 47% of all Gfi1-unique sites are associated with downregulated genes (Fig. 6E,F). In notable contrast, the majority of Gfi1 peaks with E-boxes (i.e. Gfi1-Atoh1 co-bound sites) are associated with up-regulated genes (Fig. 6D). Interestingly, these Gfi1 and Atoh1 co-bound sites do not show a preference for promoters and are more frequently located in intergenic and intronic regions (Fig. 6E). These findings support the hypothesis that Atoh1 can switch Gfi1 from a repressor to a co-activator.

We then looked for evidence for the different roles of Gfi1 by examining DE genes. We performed unsupervised clustering of all genes differentially expressed relative to no-dox controls in the iGfi1, iAtoh1, iG+A and iGPA lines. A total of 1921 genes were clustered into five groups in accordance with their shared expression patterns across these four lines. Group 4 includes a set of Atoh1-responsive genes that are repressed by the presence of Gfi1 (Supplemental Fig. S7A: Group 4). We were able to detect Gfi1 binding at the vicinity of some group 4-genes, including several Hox and Id genes, as well as TFs important for lineage differentiation (e.g. Otx1 and NeuroD4) (Supplemental Fig. S7B). This supports the suggestion that part of Gfi1’s function is the direct repression of non-HC targets. In addition, groups 1 and 2 contain genes that require Gfi1 and Atoh1 for their upregulation. Interestingly, Gfi1 enhanced the expression of common Atoh1 target genes (Supplemental Fig. S7A: Group 1-Notch and neuronal genes), as well as enabling expression of a group of genes that Atoh1 by itself is not able to activate (Supplemental Fig S7A: Group 2). Notably, this included genes associated with inner ear morphogenesis. However, the other HC-related GO term ‘sensory perception of sound’ is only associated with genes upregulated in the presence of all three TFs (Group 3), supporting the phenotypic observations that more complete iHC differentiation requires Pou4f3 in addition to Gfi1 and Atoh1.

To further test the hypothesis that Gfi1 is both a repressor and an Atoh1-dependent coactivator, we performed transcriptional assays in P19 cells transfected with reporter constructs bearing luciferase with a minimal promoter sequence under the regulation of multimerised binding sites for Gfi1 or Atoh1. We first tested the effect of Gfi1 expression on the expression of luciferase under control of a promoter that consisted of strong Gfi1 binding sites (R21). Consistent with the findings of previous studies, Gfi1 was able to repress expression of the R21 reporter gene efficiently (Zweidler-Mckay et al. l996; Grimes et al. 1996) (Fig. 6J). This corroborates our observation in iGPA cells that Gfi1 binding to its own motif is associated with transcriptional repression. We then tested a reporter gene under the control of multimerised Atoh1 E box binding sites (AtEAM)^6^. As expected, expression of Atoh1 on its own was able to induce AtEAM reporter gene expression and this effect was further enhanced by adding an E-protein (E-proteins are essential heterodimerisation partners of Atoh1) (Fig. 6H). Expression of Gfi1 alone had no effect on reporter activity consistent with inability to bind the E box, but remarkably co-expression of Gfi1 and Atoh1 significantly augmented reporter gene expression. This reveals a synergy in activity between these two TFs (Fig. 6H). Similar results were observed when Gfi1 was co-expressed with another proneural bHLH factor, Ascl1, suggesting that such synergistic effect could occur more broadly in other cell fate decisions requiring proneural TFs (Fig. 6I). Interestingly, expression of Atoh1 is also able to counteract Gfi1 repression of the R21-regulated reporter gene, presumably because Atoh1 binds directly to Gfi1. Such a derepression effect might explain why we still detect 53% of genes up-regulated when Gfi1 is bound to their regulatory regions via its DNA motifs (Fig. 6F) in the presence of Atoh1.

Taken together, these findings suggest that Gfi1 has a dual capacity to (1) act as a transcriptional repressor when bound to its DNA motifs located at promoter regions; (2) act as a transcriptional co-activator when interacting with Atoh1.

## DISCUSSION

Although the lineage-determining ability of TFs is often modulated by working in combination with other TFs, the mechanisms by which such switching occurs are not well known. In this study using a model *in vitro* system, we found that Atoh1 is repurposed from a neuronal to HC determinant by the combined activity of Gfi1 and Pou4f3. Our evidence suggests that Atoh1 repurposing entails two major mechanisms: (1) allowing Atoh1 access to genomic locations controlling the expression of sensory genes; (2) enhancing Atoh1’s activity at its HC target genes. These two events are unlocked by Pou4f3 and Gfi1, respectively, and at least part of their effect is likely by direct interaction with Atoh1 (Fig. 7).

**Figure 7.**
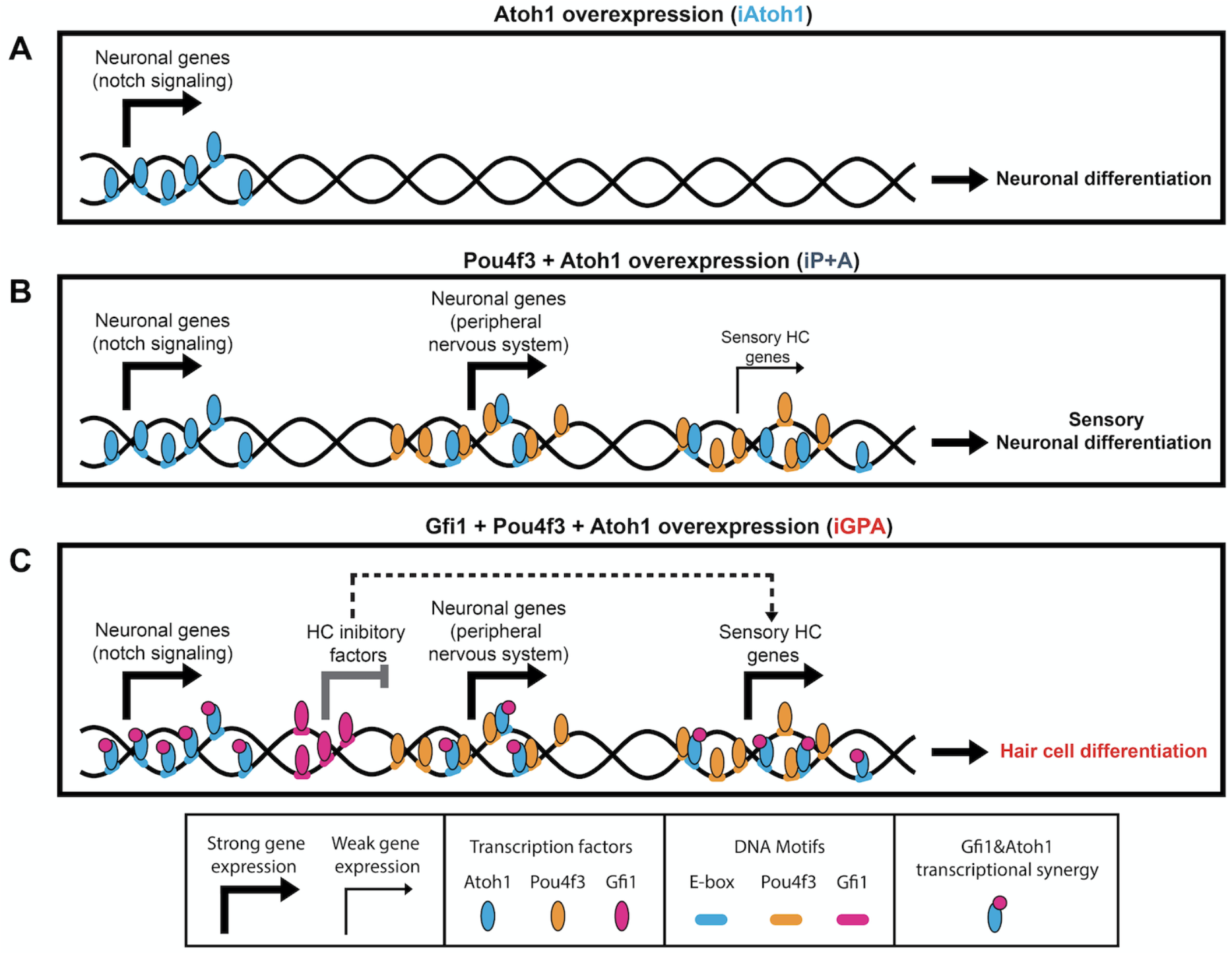
Pou4f3 and Gfi1 contribute different mechanisms in converting Atoh1 to an HC determinant in mESCs. Schematic summary of findings and possible interpretation. (A–C) Atoh1 binds to neuronal targets in all conditions. (B) Presence of Pou4f3 recruits Atoh1 (partly directly) to sensory targets (neuronal and HC) but this does not result in strong HC gene upregulation. (C) Gfi1 has a dual function in HC differentiation: it binds to and represses some targets that would otherwise interfere with HC differentiation; it cobinds via interaction with Atoh1 to sensory and HC targets where it promotes upregulation.

### Pou4f3 as a competence or pioneer factor for Atoh1

Pou4f3 expression enables the recruitment of Atoh1 to new genomic locations vital for HC differentiation. In the absence of Gfi1, however, this recruitment does not lead to full HC gene activation, rather to activation of sensory neuronal genes (Fig. 7). Given the co-occurrence of their binding motifs, part of this recruitment (at co-bound sites) is potentially by direct interaction. It is particularly striking that Pou4f3’s binding activity is unaffected upon Atoh1 (or Gfi1) expression, and yet by itself it has little capacity for transcriptional regulation in ESCs (Fig. 5C). Together, these observations suggest that Pou4f3 establishes lineage-specific competence for Atoh1-driven sensory cell differentiation, perhaps as a pioneer factor that allows Atoh1 access to its sensory targets. Consistent with this are reports that Pou domain TFs can reprogram mouse and human fibroblasts into iHC (Duran Alonso et al. 2018). Interestingly, this implies that Atoh1 is not a pioneer factor in HC differentiation *in vitro* (and perhaps *in vivo*). Indeed, *in vivo* evidence from intestinal crypt cell differentiation suggests that chromatin modification precedes Atoh1 binding (Kim et al. 2014). Together, this contrasts Atoh1 function with Ascl1, a related proneural factor that features heavily in ‘TF cocktails’ for direct reprogramming of a variety of neuronal cell lineages (Masserdotti et al. 2016). Ascl1 appears to be a pioneer factor capable of binding nucleosome occluded DNA and of promoting chromatin accessibility at its targets in neuronal differentiation (Wapinski et al. 2013; Raposo et al. 2015).

Interestingly, whilst Pou4f3 recruits Atoh1 to new targets, it does not displace Atoh1 from its default neuronal targets. Lineage switching in iGPA cells therefore does not involve complete replacement of one programme by an alternative one. This may reflect the functional similarities between HCs and neurons, e.g. both require proteins for synaptic neurotransmission.

While a major effect of Pou4f3 may be to repurpose Atoh1 directly, it should be noted that many Pou4f3 sites are distinct from Atoh1 sites, suggesting that part of its function is indirect.

### A dual function for Gfi1 as a transcriptional repressor and bHLH co-activator

In contrast to Pou4f3, Gfi1 does not alter Atoh1 binding sites. Instead Gfi1 appears to be recruited to Atoh1 sites where it acts as a co-activator of sensory genes. Thus overall, full HC determinant ability of Atoh1 requires Pou4f3 to guide it to new genomic sites, and Gfi1 to enable efficient activation of HC genes.

Our previous understanding of how Gfi1 regulates transcription largely comes from research into its role in hematopoietic development, where it acts as a major DNA-binding transcriptional repressor by recruiting chromatin regulatory complexes via its SNAG domain (McGhee et al. 2003; Duan et al. 2005; Saleque et al. 2007; Thambyrajah et al. 2016). Here we propose that Gfi1 additionally acts as a co-activator of Atoh1 target genes. Such a dual function for Gfi1 has strong support from studies of the *Drosophila* orthologue, SENS (Nolo et al. 2000) during sensory neuron specification. As well as acting as a DNA-binding transcriptional repressor, SENS promotes the activity of bHLH proneural factors, including Atonal (Acar et al. 2006; Jafar-Nejad et al. 2003b; Powell et al. 2008; 2012). In this role, SENS does not bind to its DNA motif, but directly binds proneural factors via its Zn-finger domains. Our findings suggest that such dual functionality of Gfi1/SENS is highly conserved during the specification of mechanosensory cells by Atoh1/Atonal. Gfi1 differs from SENS in possessing a SNAG domain. This might suggest that binding to Atoh1 must mask the SNAG domain to prevent its repressive activity. Interestingly, Atonal binding to SENS also interferes with its ability to repress targets (Witt et al. 2010).

In *Drosophila*, interaction of SENS with proneural factors has been proposed to constitute a temporal switch from repressor to co-activator during sensory precursor specification (Jafar-Nejad et al. 2003b). In our model, we suggest that Gfi1 repressor and co-activator roles are required concomitantly at different targets during HC differentiation. While co-activation is required for Atoh1-dependent HC gene expression, direct Gfi1-mediated gene repression is also likely to be important. A good example is the repression by Gfi1 of two bHLH TFs, Ascl1 and NeuroD4. These factors have the capacity to reprogram different cell types into neurons and so would be likely to favour Atoh1’s neurogenic activity unless repressed by Gfi1.

Future studies are needed to understand whether Gfi1 also acts as transcriptional co-activator in the hematopoietic and intestinal systems. Indeed, a few examples have been reported in which Gfi1 may activate gene expression during hematopoiesis, but in these cases Gfi1 appears to bind to its DNA motif (van der Meer et al. 2010). Atoh1/Gfi1 are co-expressed in several other cellular lineages, strongly suggesting that the synergistic mechanism may function beyond HCs. In cases where Gfi1 is not co-expressed with Atoh1 (e.g. hematopoiesis), it is possible that Gfi1 may similarly co-activate with other TFs.

### Reprogramming by survival/maintenance factors: implications for *in vitro* programming models and for HC gene therapy

*In vitro*, Gfi1 and Pou4f3 are crucial for repurposing Atoh1 for HC identity. In mouse knockout studies, however, both Gfi1 and Pou4f3 appear to be necessary for HC survival and differentiation rather than initial fate determination (Xiang et al. 1998; Wallis et al. 2003b). It is possible that the embryonic otic environment has factors and mechanisms that can compensate for these functions of Pou4f3 and Gfi1. Moreover, for Gfi1, its Atoh1-synergising role may reflect their interaction at a later stage of lineage determination/maintenance *in vivo*. It is interesting to note that this parallels SENS function in *Drosophila*: SENS appears to function during sensory lineage determination by proneural factors (Acar et al. 2006; Jafar-Nejad et al. 2003b; Powell et al. 2008) and yet SENS mutants mainly show defects in sensory lineage survival and differentiation (Nolo et al. 2000).

In this regard, the similarities between the iGPA strategy and other ‘reprogramming cocktails’ is striking (Srivastava et al. 2013; Masserdotti et al. 2016). For instance, the original ‘ABM’ regime for neuronal differentiation likewise consists of a bHLH proneural factor (Ascl1), a Pou domain TF (Brn2/Pou3f2) and Zn-finger TF (Myt1l) (Vierbuchen et al. 2010). Mytll is suggested to safeguard neuronal identity during reprogramming by repressing non-neuronal lineages (Mall et al. 2017). Gfi1 may be acting similarly to repress inappropriate gene expression during HC differentiation.

The mechanisms by which Gfi1, Pou4f3 and Atoh1 orchestrate a reprogramming event towards HC fate may have implications for developing new therapies. HC loss is a major cause of human sensorineural hearing loss, which is permanent because HCs do not regenerate in mammals. There is much interest in inducing HC regeneration using vector-supplied *Atoh1* (Richardson and Atkinson 2015; Zhang et al. 2018). Rodent models demonstrate that cochlear overexpression of Atoh1 is sufficient to generate new ectopic HCs during embryonic inner ear development, but it fails at later postnatal/adult stages (Liu et al. 2012). Our findings that Pou4f3 facilitates Atoh1 binding to HC genes and Gfi1 efficiently enhances Atoh1 activation of HC genes suggest several avenues for enhancing the therapeutic potential of Atoh1.

## MATERIALS AND METHODS

### Generation, growth and differentiation of doxycycline-inducible mESC lines

The construction procedures for the plasmids containing polycistronic sequences required to engineer all dox-inducible mESC lines used in this study are described in Supplemental Methods. These plasmids were nucleofected (Amaxa 4D-Nucleofector, Lonza) into A2Lox cells (gift from Prof. Lesley Forrester) as described previously (Iacovino et al. 2011; 2014) Cells were subsequently plated on neomycin-resistant, gamma irradiated MEF feeder cells (Stem Cell Technology, #00323). Two days after nucleofection, the recombined cells were selected by using DMEM medium supplemented with 350 μg/ml G418 (InvivoGen). Individual colonies were picked after 1 week (Iacovino et al. 2014). Medium composition, growth and differentiation protocols used for all dox-inducible mESC lines are documented in detail in Supplemental Methods.

### Immunocytochemistry and Imaging

EBs (6, 8 and 12 days old) were fixed with 1% paraformaldehyde during 15 min at room temperature (RT). Fixed EBs were cryoprotected in 15% sucrose in PBS, embedded in a solution containing 7.5% gelatine (Sigma-Aldrich) and 15% sucrose in PBS, frozen and cryosectioned (8-10 μm). EB sections were immersed in PBS at 37C until gelatin was completely dissolved, and then processed for immunocytochemistry. Sections were blocked with 10% FBS and 0.05% Tween in PBS for 1 hour, followed by incubation overnight with the following primary antibodies: Anti-MyoVIIa (1:400, HPA028918, Sigma), Anti-MyoVI (1:50, 25-6791, Proteus Biosciences) Anti-Pou4f3 (1:50, HPA038215, Sigma), Anti-Espin (1:1000, gift of A. J. Hudspeth), Anti-Gfi1 (1:2000, gift of Hugo Bellen), Anti-Gfi1 (ab21061 or ab290, Abcam), Anti-Lhx3 (1:100, ab14555, Abcam), Anti-Tuj1 (1:500, MMS-435P, Covance), Anti-GFP (1:500, ab13970, Abcam), Anti-Doublecortin (1:1000, AB2253, Merck Millipore). Sections were washed 3 times in PBS followed by incubation for 1 hour at RT with AlexaFluor-conjugated secondary antibodies (1:400, Molecular Probes) and 0.15% DAPI (Sigma-Aldrich). Slides where then mounted with prolong gold (Life technologies). Fluorescent images of fixed sections were captured with Widefield Zeiss observer or using a Leica TCS SP8 Confocal 4 Detectors. All digital images were formatted with Adobe Photoshop CS and ImageJ.

### Flow cytometry

For live cell analyses to measure Venus expression in all dox-inducible lines, EBs were dissociated and re-suspended in PBS with 2% FBS after 48h, 4 and 8 days of dox treatment. All cells were analysed FACSCalibur (BD Biosciences). Data were analysed with FlowJo software (Tree Star).

### RNA extraction and real-time quantitative PCR

EBs (6, 8 and 12 days old) were dissociated and 106 cells were used to extract total RNA following manufacturer’s instructions of the Absolutely RNA Miniprep Kit (#400800, Agilent Technologies). cDNA was synthesised with 200 ng-1 μg of total RNA using M-MLV Reverse Transcriptase (Invitrogen) and random hexamers. Quantitative PCR was performed using Light Cycler 480 SYBR Green I Master Mix (Roche) in a LightCycler 480 II Instrument. Primer sequences are described in Supplemental Methods.

### RNA sequencing (RNA-seq)

Total RNA was extracted from untreated EBs at day 6, and from FACS sorted EBs at day 6 previously treated with Dox for 48h. Cell sorting of Venus+ populations were done on a FACS Aria cell sorter (Becton Dickinson). RNA concentration and purity was determined by spectrophotometry and integrity was confirmed using a High Sensitivity RNA ScreenTape system on an Agilent 2200 TapeStation. RNA-seq libraries were generated with 1ug of total RNA using KAPA mRNA HyperPrep Kit (KK8580, Kapa Biosystems) following manufacturer’s instructions. RNA-seq libraries were pooled at equal concentration and sequenced with Edinburgh Genomics on the Illumina HiSeq 4000 platform. Data analyses for the RNA-seq experiments are described in Supplemental Methods.

### Chromatin Immunoprecipitation followed by sequencing (ChIP-seq)

ChIP-seq experiments were performed by adapting and modifying protocols described in (Ruetz et al. 2017; Ford et al. 2014). Further details on ChIP-seq procedures and data analyses are described in Supplemental Methods.

### GST Pulldown Assay and Luciferase reporter assay

The GST Pulldown assay was carried out as described by (Nguyen and Goodrich 2006) with some modifications which are documented in Supplemental Methods. The procedures for luciferase reporter experiments can be also found in Supplemental Methods including the plasmids used for both assays.

### Statistics

All data are expressed as mean±s.e.m. and statistical significance was assessed using an unpaired Student’s t-test. For all statistics, data from at least three biologically independent experiments were used. Data and graphs were tabulated and prepared using Microsoft Excel and GraphPad Prism software. P<0.05 was considered statistically significant.

### Data availability

RNA-seq and ChIP-seq data are available through Gene Expression Omnibus under accession number GSE136909.

## ACKNOWLEDGEMENTS

We are very grateful to James Ashmore and other members of the Tomlinson group for expert advice in bioinformatics, and to Tyson Reutz and members of the Kaji and Chambers labs for advice on ChiP Seq experiments. We thank Ian Simpson for his computational help in the early stages of the project. We also thank Andy Groves, Stephanie Juniat, Nicolas Daudet and Jonathan Gale for helpful discussions, and also Luis Sanchez-Guardado and Domingos Henrique for support during the early stages of this project. We thank Steve Pollard, Ian Chambers and members of the Jarman and Lowell labs for technical advice and comments on the manuscript. This work was supported by a Marie Skłodowska-Curie Individual Fellowship to A.C. and A.P.J. (707015 — GRNHairCell), a BBSCR/UKRI Flexible Talent Mobility Award to A.C., Wellcome Trust Senior Fellowship (WT103789AIA) to S.L., MRC project grant (MR/L021099/1) to S.L. and A.P.J, an MRC career development award (MR/N024028/1) and by UK Research Councils’ Synthetic Biology for Growth programme of the BBSRC, EPSRC and MRC (BB/M0180404/1) to A.S.

## Author contributions

A.C., S.L. and A.P.J. designed the bulk of the study, with contributions from L.M.P. and A.S. A.C. performed all the experiments except for the luciferase assays and pull-down, which were performed by L.M.P. All authors critically evaluated the data. A.C., S.L. and A.P.J. wrote the paper with input from all authors.

