## Supplemental figures and methods for "Atoh1 is repurposed from neuronal to hair cell determinant by Gfi1 acting as a coactivator without redistributing Atoh1’s genomic binding sites"

Costa_FigS1

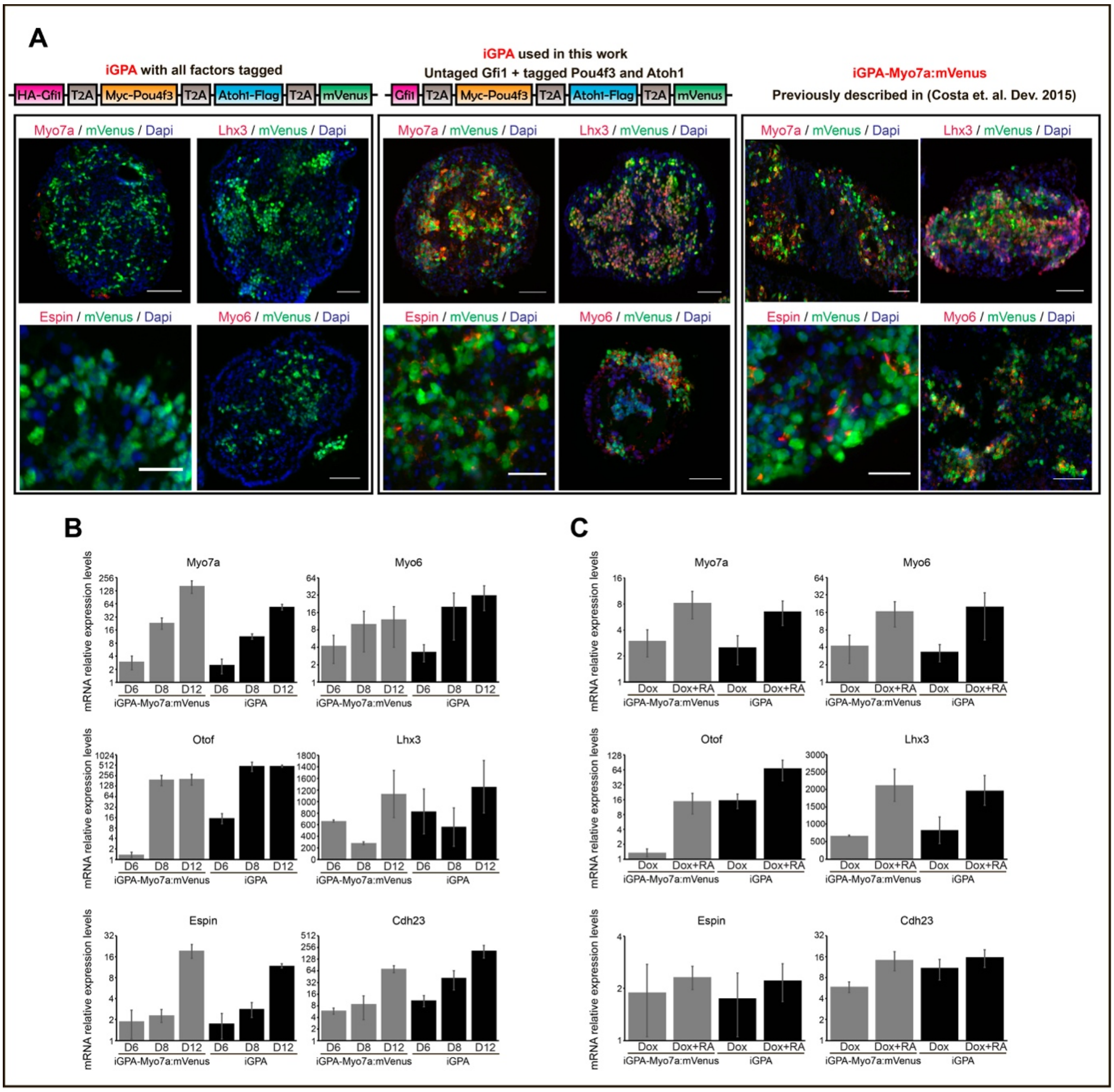


**Figure S1. A double-tagged iGPA cell line supports HC differentiation.** (A) Above: schematics of constructs used (triple-tagged line, double-tagged line (‘iGPA’ in this work), original untagged line (‘iGPA-Myo7a:mVenus’). Below: expression of HC markers (red) in induced EBs relative to mVenus (green) and DAPI (blue). Double tagged-line supports HC marker expression but triple-tagged line does not. (B) qPCR in EBs for expression of HC markers. Time course after induction for iGPA-Myo7a:mVenus’ (grey) and double-tagged iGPA (black) cell lines. (C) qPCR in EBs for expression of HC markers. Addition of RA increases expression of HC markers in the double-tagged iGPA line.

A

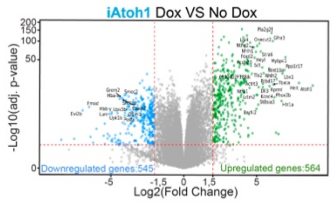


GO analysis of 564 upregulated genes

| Go term (biological process) | P-value | Genes |
|---|---|---|
| Neuron migration | $1.1\times10^{-8}$ | Barhl2, Prrxl1, Nr2f2, Arx, Astn1, Atoh1, Neurod4, Cck, Cdk5r1, Cdk5r2, Dcc, Dcx, Gfra3, Mnx1, Ascl1, Phox2b, Unc5d, Satb2, Vax1, Barhl1, Mdga1 |
| Compartment pattern specification | $1.0\times10^{-6}$ | Dll1, Dll3, Lfng, Notch1 |
| Positive regulation of Notch signaling pathway | $1.4\times10^{-6}$ | Dll1, Hes5, Jag1, Jag2, Lfng, Ascl1, Mfng, Notch1, Dll4 |
| Regulation of somitogenesis | $1.7\times10^{-6}$ | Dll1, Lfng, Notch1, Pax3, Dmrt2, Hes7 |
| Axon guidance | $9.6\times10^{-6}$ | Unc5a, Prrxl1, Arx, Atoh1, Cdk5r1, Dcc, Epha8, Gfra3, Mnx1, Lhx3, Ngfr, Ntrk1, Sema6c, Slit1, Unc5d, Tubb3, Vax1, Rnf165, Robo2, Nfasc, Ephb1, Nrcam, Draxin |
| Negative regulation of osteoblast differentiation | $2.2\times10^{-5}$ | Areg, Chrd, Cited1, Gdf10, Id1, Id2, Id3, Notch1, Mir135a-1 |
| Noradrenergic neuron development | $3.3\times10^{-5}$ | Ascl1, Phox2b, Insm1 |
| Embryonic skeletal system morphogenesis | $3.8\times10^{-5}$ | Dscaml1, Runx2, Gsc, Hoxa3, Hoxa5, Hoxa7, Hoxb3, Hoxd10, Hoxd11, Prrx2, Satb2, Tbx1, Tgfb3 |
| Nervous system development | $7.2\times10^{-5}$ | Barhl2, Unc5a, Prrxl1, Syt17, Chrna3, Aatk, Foxp2, Asic2, Ace, Dscaml1, Vsx1, Adm, Nr2f2, Areg, Arx, Astn1, Atoh1, Neurod4, Bche, Fabp7, Camk2a, Runx2, Cck, Cdk5r1, Cdk5r2, Chrd, Cited1, Cpne6, Dcc, Dcx, Dll1, Dll3, Dmd, Epha8, Fgf14, Gal, Gfra2, Gfra3, Gh, Gnat1, Gsc, Hes5, Mnx1, Hmx3, Hoxa3, Hoxb3, Hoxd10, Hprt, Elavl3, Elavl4, Id1, Id2, Id3, Jag1, Jag2, Ush1g, Lbx1, Lhx3, Mycl, Lrrn1, Lrrn2, Ntng2, Ascl1, Foxc1, Map6, Myoc, Myt1, Nbl1, Neurod1, Nefl, Ngfr, Notch1, Nrl, Ntrk1, Otx1, Pak3, Pax3, Pitx3, Phox2b, Pou3f1, Pou3f2, Rxrg, Sct, Sema6b, Sema6c, Slc1a2, Slit1, Snap25, Sox10, Sall3, Wnt8a, Unc5d, Lzts1, Lrtm2, Satb2, Tbx1, Tlx2, Tubb3, Uncx, Vax1, Wnt10a, Wnt10b, Onecut2, Rnf165, Rnf112, Clrn1, Prdm13, Helt, Sh2b2, Mgll, Tenm1, Brinp2, Zfp804a, Lrrc38, Lgi4, Myo16, Dusp15, Fev, Adgre5, Dpysl4, Robo2, Nfasc, Rtn4rl2, Ephb1, Lrrc4b, Zbtb18, Nrcam, Islr2, Lingo4, Foxo6, Miat, Rbfox3, Cntnap1, Insm1, Barhl1, Dll4, Ccdc141, Hes6, Heyl, Ntn4, Bhlhe22, Gprc5b, Rtn4r, Nrn1, Srrm4, Draxin, Ttll7, Atoh8, Faim2, Whrn, Mdga1, Mmd2, Slitrk5, Prdm8, Kcnip2, Igsf9 |

B

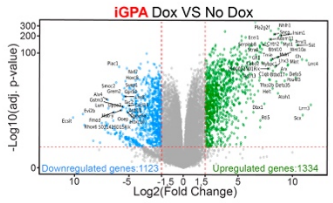


GO analysis of 1334 upregulated genes

| Go term (biological process) | P-value | Genes |
|---|---|---|
| Locomotory behavior | $1.1\times10^{-8}$ | Chrnb4, Grm6, Chrna3, Adora2a, Adra1b, Alk, Apba2, Astn1, Atp2b2, Pcdh15, Cacna1b, Calb1, Celsr1, Chat, Chl1, Cnp, Crhr1, Slc6a3, Dnm1, Drd4, Dmbx1, Fgf14, Hprt, Elavl4, Kcnd2, Lmx1b, Myo15, Myo6, Npy1r, Nr4a2, Selenop, Sncg, Nrg1, Cntn2, Slc18a2, Th, Tshr, Adcy5, Slitrk6, Pak7, Lgi4, Olfm2, Adgrl3, Apba1, Usp2, Espn, Sez6l, Gprc5b, Gpr88, Rasd2, Atp1a2 |
| Regulation of ion transmembrane transport | $1.0\times10^{-6}$ | Pcsk9, Scn4a, Fxyd2, Cacna1b, Crhr1, Drd4, Cacng5, Rem2, Fgf14, Gal, Hpca, Hrc, Htr3a, Kcna4, Kcna7, Kcnab2, Kcnb1, Kcnc1, Kcnc3, Kcnd2, Kcnh3, Kcnj11, Kcnj12, Kcnj2, Kcnj3, Kcnj5, Kcnj8, Kcnj9, Kcnmb1, Fxyd3, Nos1, Bpifa1, Ppargc1a, Tmc2, Ptk2b, Pirt, S100a1, Scn10a, Scn5a, Abcc8, Cnksr3, Acsl6, Kcnk13, Thy1, Clic5, Amigo1, Osr1, Kcng1, Lrrc38, Mtnr1b, Kcne2, Gsg1l, Cacng4, Cacng6, Cacna1f, Fxyd1, Rrad, Kcnip3, Cacna2d2, Trpm5, Fxyd7, Tesc, Cacna1h, Kcnq4, Gpr35, Tspan13, Kcne1l, Kcnv1, Kcnip1, Jsrp1, Shisa9, Cnih3, Wnk2, Nol3, Atp1a2, Kcnb2, Kcnc4 |
| Sensory perception of sound | $1.4\times10^{-6}$ | Atp2b2, Pcdh15, Col11a2, Col2a1, Crym, Dnm1, Bsnd, Eya4, Hpn, Lrig1, Jag2, Ush1g, Map1a, Myo15, Myo6, Pou4f2, Pou4f3, Tmc2, Ripor2, Tmie, Tbx1, Slc17a8, Ush2a, Clic5, Slitrk6, Loxhd1, Otoa, Myo3b, Grxcr2, Barhl1, Espn, Kcnq4, Mir124-2hg, Ush1c, Tomt, Otof |
| Neuron migration | $1.7\times10^{-6}$ | Barhl2, Prrxl1, Nr2f2, Arx, Astn1, Atoh1, Neurod4, Cck, Cdk5r2, Celsr1, Chl1, Dcc, Dcx, Erbb4, Gfra3, Mnx1, Lhx1, Lmx1b, Scrt1, Met, Ntrk3, Nr4a2, Ptprz1, Sox1, Sox14, Spock1, Unc5d, Nrg1, Satb2, Cntn2, Lrp12, Disc1, Tlx3, Adgrl3, Barhl1, Olfm1, Ndnf, Mdga1, Olig3 |
| Homophilic cell adhesion via plasma memb... | $9.6\times10^{-6}$ | Cdh22, Dscaml1, Pcdh15, Cdh15, Cdh4, Celsr1, L1cam, Pcdh10, Pcdh8, Pcdh20, Amigo1, Cdh20, Cdh7, Cadm4, Robo2, Cdh26, Cadm1, Clstn2, Cdhr3, Cdhr5, Pcdhb18 |
| Inner ear morphogenesis | $2.2\times10^{-5}$ | Atoh1, Atp2b2, Pcdh15, Celsr1, Col2a1, Dll1, Eya1, Fgf9, Gbx2, Gfi1, Hmx3, Hpn, Lrig1, Ush1g, Myo15, Myo6, Neurog1, Pou4f3, Ripor2, Tmie, Tbx1, Stox1, Tshr, Clic5, Slitrk6, Myo3b, Kcnq4, Ush1c |
| Potassium ion transmembrane transport | $3.3\times10^{-5}$ | Gal, Hpn, Kcna4, Kcna7, Kcnab2, Kcnb1, Kcnc1, Kcnc3, Kcnd2, Kcnh3, Kcnj11, Kcnj12, Kcnj2, Kcnj3, Kcnj5, Kcnj8, Kcnj9, Kcnmb1, Abcc8, Abcc9, Kcnk12, Kcnk13, Kcnk9, Slc9a2, Amigo1, Kcng1, Lrrc38, Mtnr1b, Kcne2, Kcnip3, Kcnq4, Kcne1l, Kcnv1, Kcnip1, Tmem38a, Atp1a2, Kcnb2, Kcnc4 |
| Axon guidance | $3.8\times10^{-5}$ | Unc5a, Prrxl1, Arx, Atoh1, Bmp7, Cdh4, Chl1, Dcc, Efna3, Epha8, Gbx2, Gfra3, Mnx1, Isl1, L1cam, Lhx1, Lhx3, Ngfr, Ntrk1, Pou4f2, Pou4f3, Ptpro, Sema3c, Sema3f, Sema6c, Slit1, Unc5d, Cntn2, Tbr1, Zic2, Robo2, Nfasc, Nrcam, Lgi1, Bcl11b, Dpysl5, Draxin |
| Axonogenesis | $7.2\times10^{-5}$ | Barhl2, Unc5a, Prrxl1, Aatk, Arx, Atoh1, Bmp7, Cck, Cdh4, Chl1, Cnp, Dcc, Dcx, Efna3, Epha8, Gbx2, Gfra3, Mnx1, Isl1, L1cam, Lhx1, Lhx3, Lrrn2, Ntng2, Map2, Myo5b, Nefl, Nefm, Ngfr, Nin, Nptx1, Ntrk1, Ntrk3, Nr4a2, Pou4f2, Pou4f3, Ptpro, Ptprz1, Rab3a, Sema3c, Sema3f, Sema6c, Sema7a, Shox2, Slit1, Unc5d, Nrg1, Cntn2, Tbr1, Thy1, Zic2, Amigo1, Plppr4, Lingo1, Slitrk6, Mgll, Twf2, Lrrc38, Disc1, Robo2, Nfasc, Nrcam, Fstl4, Islr2, Lingo4, Adcy1, Cacna1f, Olfm1, Lgi1, Bcl11b, Rtn4r, Lpar3, Dpysl5, Plxnd1, Nrn1, Draxin, Brsk2 |

C

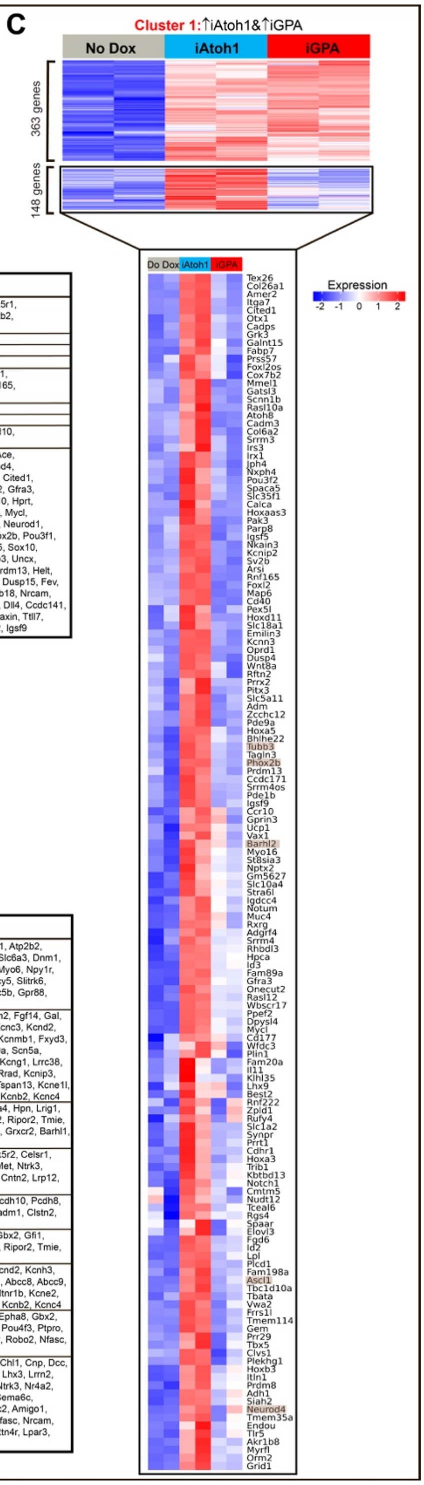


**Figure S2. Transcriptomes from iGPA and iAtoh1 EBs.** (A) Volcano plot of iAtoh1 EBs (Dox induced vs uninduced); GO analysis of upregulated genes showing nine most significant terms. (B) Volcano plot of iGPA EBs (Dox induced vs uninduced); GO analysis of upregulated genes. (C) Cluster analysis of genes upregulated in iAtoh1 cells (cluster 1), divided according to presence or absence of expression in iGPA cells (clusters 1A and 1B). The genes in cluster 1B (genes expressed in iAtoh1 but not iGPA) are shown expanded below.

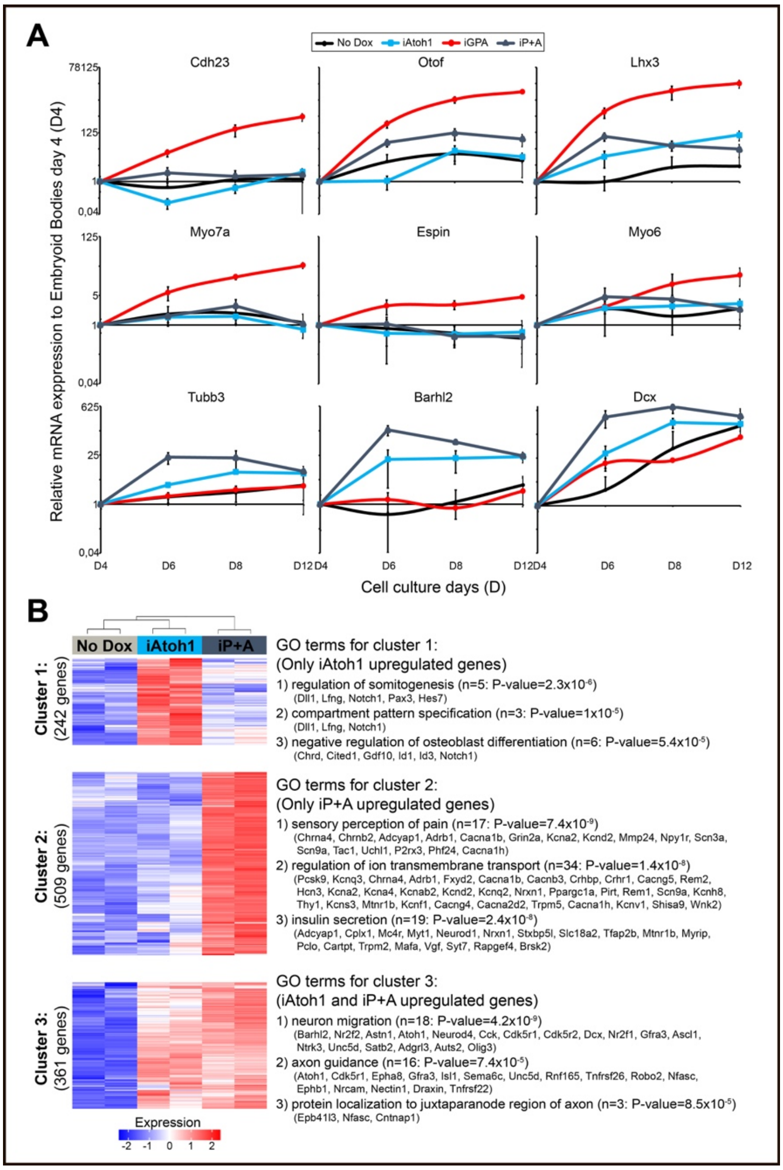


**Figure S3. Coexpression of Atoh1 and Pou4f3 (iP+A cell line) induces sensory but not HC genes.** (A) qPCR in EBs of iAto1, iGPA and iP+A for HC (Cdh23, Otof, Lhx3, Myo7a, Espin, Myo6) and neuronal (Tubb3, Barhl2, Dcx) markers. (B) Cluster analysis of transcriptomes for iAtoh1 and iP+A EBs.

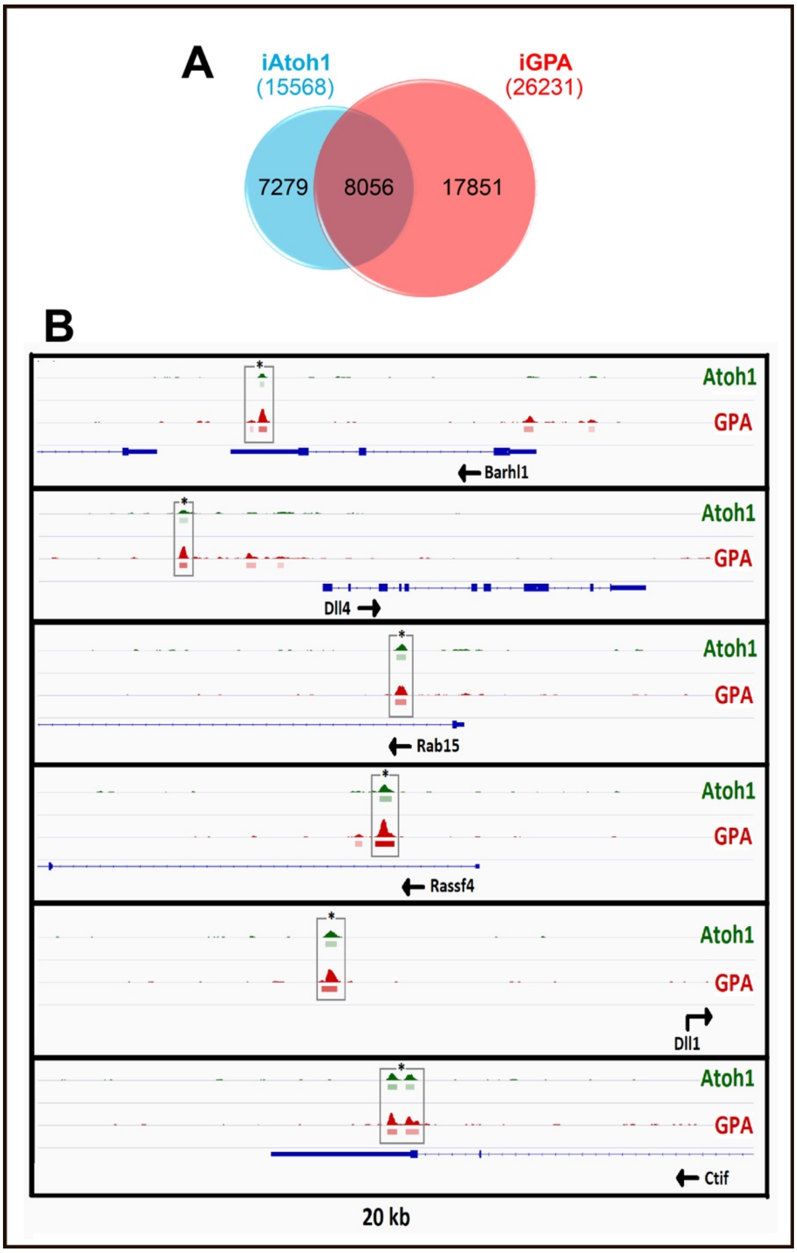


**Figure S4. Atoh1 genomic binding sites in iAtoh1 and iGPA cells.** (A) Venn diagram of Atoh1 genomic binding sites in iAtoh1 and iGPA cells. (B) Atoh1 target genes shared between iAtoh1 and iGPA cells. Examples of genes differentially expressed in both iAtoh1 and iGPA cells that also share Atoh1 binding peaks.

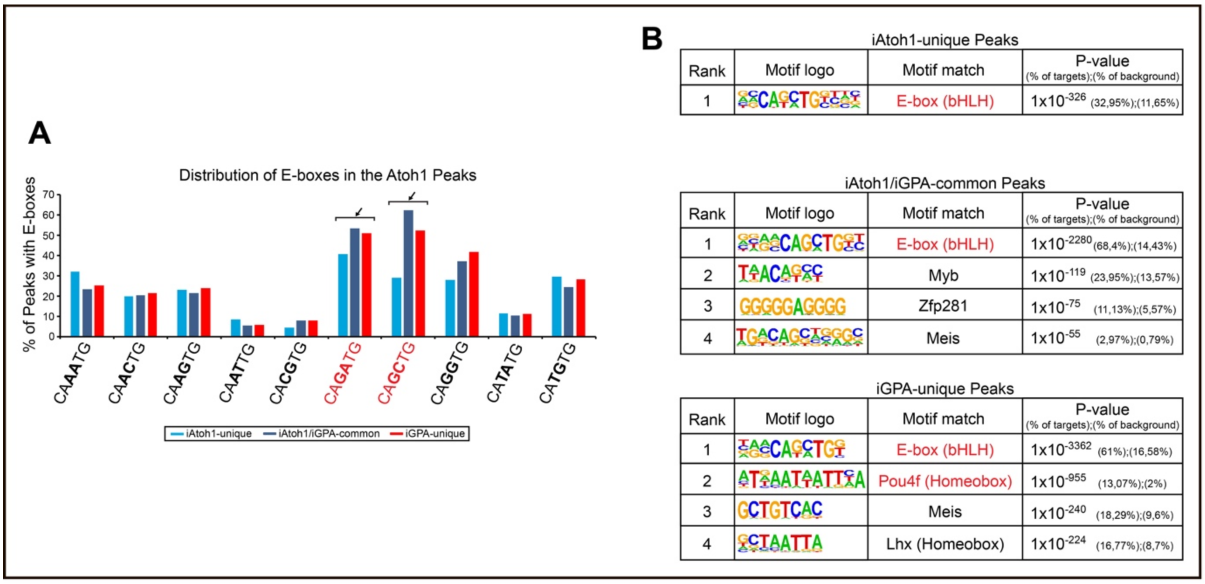


**Figure S5. DNA motifs in Atoh1 binding peaks.** (A) Number of E box motifs in binding peaks associated with the three classes of Atoh1 binding site in Fig. 4A. (B) *de novo* sequence motifs enriched in the three classes of Atoh1 binding peaks. E box motifs are detected in all classes and Pou4f motifs are detected in the iGPA-unique peaks. Gfi1 motifs are not detected at all.

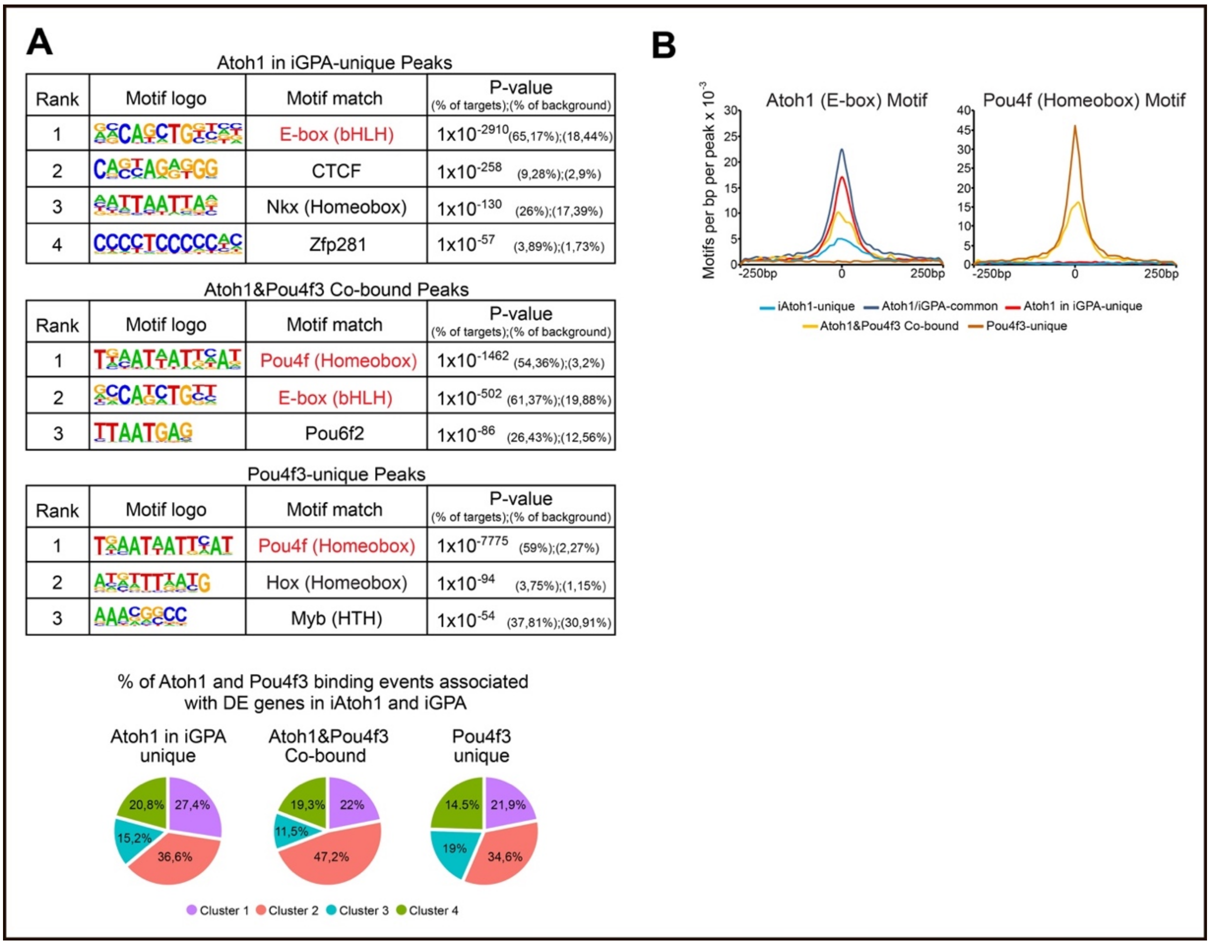


**Figure S6. Direct binding of Atoh1 and Pou4f3 to a subset of genomic locations.** (A) *de novo* sequence motifs enriched in Atoh1/Pou4f3 co-bound peaks from iGPA cells (middle) compared with peaks bound only by Atoh1 (upper) or Pou4f3 (lower). (B) Atoh1 and Pou4f DNA motifs associated with Atoh1 and Pou4f3 binding peaks.

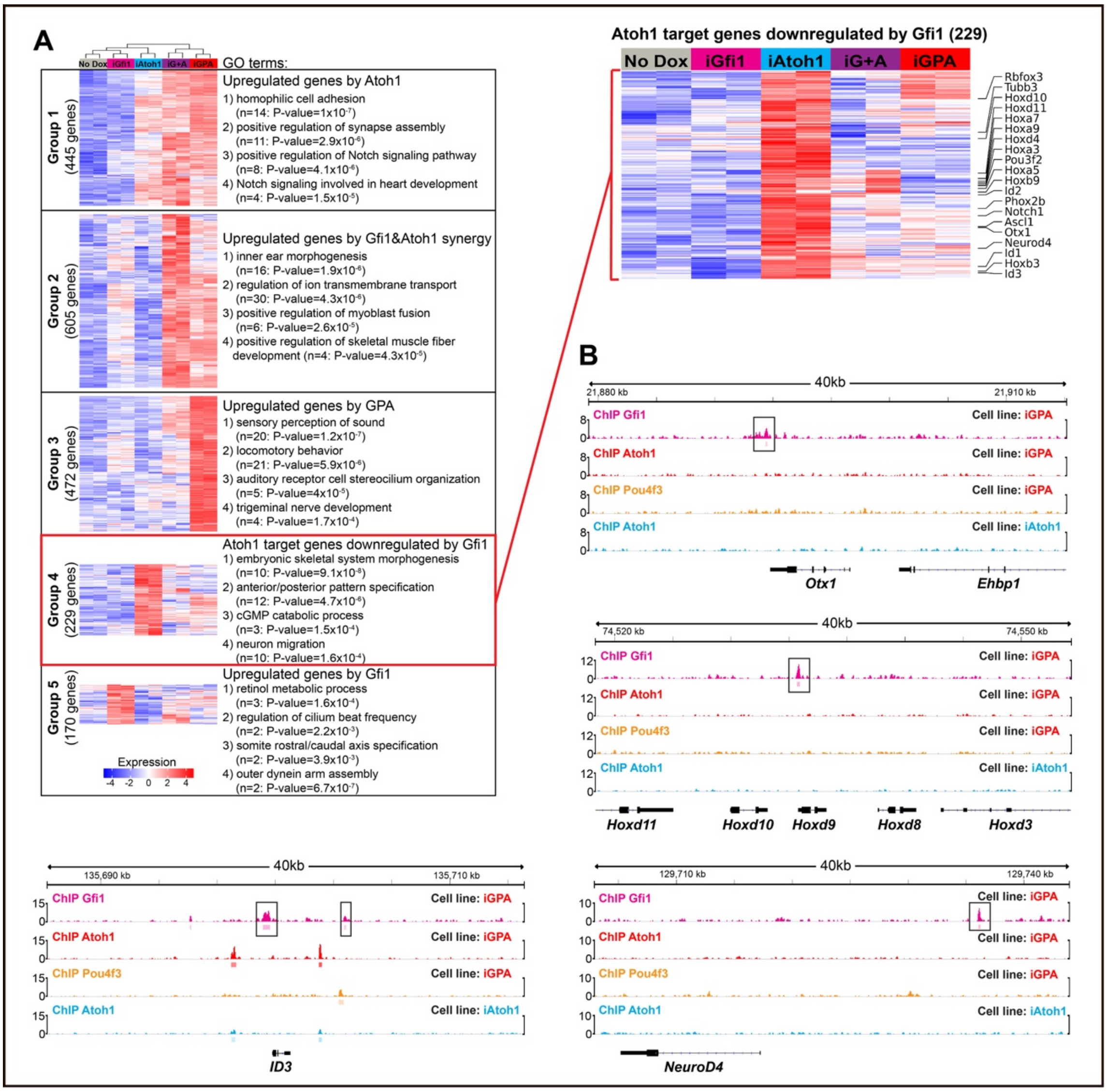

**Figure S7. Gfi1 has both positive and negative effects on Atoh1-induced transcriptomes.** (A) Cluster analysis of gene expression in iGfi1, iAtoh1, iG+A and iGPA cells. Group 4 (Atoh1-dependent genes that are repressed by presence of Gfi1) is shown expanded to the right. (B) Examples of Gfi1 binding peaks at Gfi1-repressed genes.

### Supplemental Methods

#### Plasmids construction for Dox-inducible mESCs lines:

For the generation of the iAtoh1, iPou4f3, iGfi1, iG+P, iG+A, iP+A and iGPA dox-inducible mESC lines the following plasmid were constructed following the procedures described below. All plasmids and full sequences are available upon request:

p2Lox-Atoh1F-Venus: A 3xFlag tag was added to the 3' end of the murine Atoh1 coding sequence by PCR amplification using the primers (Fwd_EcoRI_XhoI_Atoh1: TCGGAATTCCTCGAGGCCACCATGTCCCGCCTGCTGCAT and REV_Atoh1_3XFlag_XbaI: GCATCTAGACTTGTCATCGTCATCCTTGTAATCGATATCATGATCTTTATAATCACCGTCATGGTCTTTGTAGTC ACTGGCCTCATCAGAGTCACTG) and the GPAPlox vector (Costa et al. 2015) as a template. The PCR product was cloned into a 2AP-mVenusNLSPlox vector (Costa et al. 2015 unpublished plasmid). Next, the entire sequence the Atoh1-3xFlag-2AP-mVenusNLS was excised and cloned into the p2lox plasmid (Addgene #34635) (Iacovino et al. 2011).

p2Lox-MPou4f3-Venus: A 3xMyc was added to the 5' end of the murine Pou4f3 coding sequence by PCR amplification using the primers (Fwd_EcoRI_MluI_3XMyc_Pou4f3: TCGGAATTCACGCGTGCCACCATGGAGCAGAAGCTGATCTCCGAGGAGGACCTGAACGAACAAAAACTCATC TCAGAAGAGGATCTGAACGAGCAGAAGCTGATCTCCGAGGAGGACCTGATGATGGCCATGAACGCC and Rev_Pou4f3_XbaI: CGATCTAGAGTGGACAGCAGAGTATTTCATTCG) and the GPAPlox as a template. The PCR product was cloned into a 2AP-mVenusNLSPlox vector and them, the entire sequence the 3xMyc-Pou4f3-2AP-mVenusNLS was excised and cloned into the p2lox.

p2Lox-Gfi1-Venus: Murine *Gfi1* ORF was PCR amplified from the GPAPlox plasmid using the primers (Fwd_XhoI_Gfi1: ATAAGCTCGAGGCCACCATGCCGCGCTC and Rev_Gfi1_XbaI: CGATCTAGATTTGAGTCCATGCTGAGTCTCTCG). The PCR product was cloned into a 2AP-mVenusNLSPlox vector and them, the entire sequence the Gfi1-2AP-mVenusNLS was excised and cloned into the p2lox.

p2Lox-Gfi1+MPou4f3-Venus: Murine *Gfi1* ORF was PCR amplified from the GPAPlox plasmid using the primers (Fwd_XhoI_Gfi1: ATAAGCTCGAGGCCACCATGCCGCGCTC and Rev_2APGfi1_MluI: CGAACGCGTAGGGCCGGGGTTCTCCTC). The PCR product was cloned into the p2Lox-MPou4f3-Venus plasmid.

p2Lox-Gfi1+Atoh1F-Venus: The 3xFlag tag Atoh1 sequence was PCR amplified using p2Lox-Atoh1F-Venus as a template and the following primers (Fwd_MluI_Atoh1: ATAACGCGTATGTCCCGCCTGCTGCAT and Rev _Xbla3xFlag_Atoh1: GCATCTAGACTTGTCATCGTCATCCTTGT). This PCR product was inserted into the p2Lox-Gfi1+MPou4f3-Venus plasmid by excising first the 3xMyc-Pou4f3 coding sequence and replacing it with 3xFlag-Atoh1.

p2Lox-MPou4f3+Atoh1F-Venus: The 3xMyc tag was added to the 5' end of the Pou4f3 coding sequence by PCR amplification using the primers (Fwd_EcoRI_MluI_3XMyc_Pou4f3: TCGGAATTCACGCGTGCCACCATGGAGCAGAAGCTGATCTCCGAGGAGGACCTGAACGAACAAAAACTCATC TCAGAAGAGGATCTGAACGAGCAGAAGCTGATCTCCGAGGAGGACCTGATGATGGCCATGAACGCC and Rev_Pou4f3_BamHI: CGAGGATCCGTGGACAGCAGAGTATTTCATTCG) and the GPAPlox as a template. The PCR product was cloned into the Atoh1-3xFlag-2AP-mVenusNLSPlox vector and the entire sequence the Myc3x-Pou4f3-2AP-Atoh1-3xFlag-2AP-mVenusNLS was excised and cloned into the p2lox.

p2Lox-Gfi1+MPou4f3+Atoh1F-Venus: *Gfi1* ORF was PCR amplified from the GPAPlox plasmid using the primers Fwd_XhoI_Gfi1: ATAAGCTCGAGGCCACCATGCCGCGCTC and Rev_2APGfi1_MluI: CGAACGCGTAGGGCCGGGGTTCTCCTC). The PCR product was cloned into the Myc3x-Pou4f3-2AP-Atoh1-3xFlag-2AP-mVenusNLSPlox vector and the entire sequence the Gfi1-2AP-Myc3x-Pou4f3-2AP-Atoh1-3xFlag-2AP-mVenusNLS was excised and cloned into the p2lox.

### Plasmids construction for GST Pulldown and Luciferase reporter assays

The GST-Gfi1 bacterial expression construct comprising the mouse Gfi1 coding sequence cloned in pGEX-KG between Bam HI and EcoRI sites was a kind gift from Dr Angela Chen and Dr Pin Yao Wang, Taiwan. The pGBKT7-myc-Atoh1 construct was made by ligating PCR amplified mouse Atoh1 coding sequence in pGBKT7 between EcoRI and NdeI.

Mouse Gfi1 expression constructs used in P19 cells included pEx-Mm02706-M12 (Genecopoeia) and pCMV5-Gfi1 which was made by PCR amplification of the Gfi1 coding sequence and cloning in pCMV5 between MluI and XbaI. pCMV-myc-Atoh1 was made by amplification of the Atoh1 coding sequence and cloning in pCMV-myc-N (Clontech) between EcoRI and XhoI.

The Ascl1 expression construct pCDNA Mash1-HA, the hE47 expression construct pRC-CMV-hE47, and the Ascl1-specific E-box luciferase reporter construct pGL3-6*AsclE1 were kind donations from Diogo Castro (Castro et al. 2006).

Primers to amplify Atoh1 to make the pGBKT7-myc-Atoh1 expression construct: Atoh1 was amplified from mouse genomic DNA by PCR using the primers below. The PCR product was digested with EcoRI and NdeI then ligated in pGBKT7 (Clontech) digested with the same restriction enzymes and treated with Antarctic phosphatase. Atoh1-NdeI forward (GCGCATATGTCCCGCCTGCTGCATGCAG), Atoh1-EcoRI reverse (GACCGGAATTCCTAACTGGCCTCATCAGAGT)

Primers for production of E-box concatemers used in luciferase reporter constructs: Concatemers were made for the following as described in (Powell et al. 2004), and cloned in either pGL4.23 luciferase (AtEAM) or pGL3-promoter luciferase (R21-Gfi1 motif) reporter vectors (Promega).

Primers for Atoh1 E-box Associated Motif construct (pGL4.23-6*AtEAM) with E-box sequence according to (Klisch et al. 2011): AtEAM forward (with BglII for multimerisation: GATCT GGAGCGCCAGCTGGCAGGTTGGAGCGCCAGCTGGCAGGTTG), AtEAM reverse (with BamHI for multimerisation: GATCCAACCTGCCAGCTGGCGCTCC AACCTGCCAGCTGGCGCTCCA).

Primers for R21 Gfi1 binding site luciferase reporter construct (pGL3-6*R21): R21 forward (GATCTCACATAAATCACTGCCCACATAAATCACTGCCG), R21 reverse (GATCC GGCAGTGATTTATGTGGGCAGTGATTTATGTGA

Primers to amplify Gfi1 sequence to make pCMV5-Gfi1 expression construct: MluI-Gfi1 forward (GCGACGCGTGCCACCATGCCGCGCTCATTCCTG), Gfi1-STOP-XbaI reverse (GCGTCTAGACTATTTGAGTCCATGCTGAGTCTCTCGGTGCCTCC)

Primers to amplify Atoh1 sequence to make pCMV-myc-Atoh1 construct: Atoh1-EcoRI forward (GCGAATTCGGATGTCCCGCCTGCTGCATGCAG), Atoh1-XhoI reverse (GCGCTCGAGCTAACTGGCCTCATCAGAGT).

### mESC maintenance and embryoid body (EB) differentiation

mESCs were routinely grown on top of mitotically inactivated mouse embryonic fibroblasts (MEFs) at 37C in a 5% CO2 incubator in Dulbecco's Modified Eagle's Medium (DMEM), supplemented with 10% fetal bovine serum (FBS) (ES-qualified), 100U/ml human leukemia inhibitory factor (LIF), 2 mM L-Glutamine, 1 mM sodium pyruvate, 1 mM 2-mercaptoethanol,1% of penicillin-streptomycin and 1x non-essential amino acids (all from Life Technologies). Cells were passaged every other day, at constant plating density of 3×104 cells/cm2.

For EB differentiation, MEFs were first depleted from the mESCs by using 0.25% trypsin-EDTA (Life Technologies) to dissociate MEFs-coated mESCs culture dishes into a single cell suspension which was incubated on gelatin-coated (0.1%) dishes for 45 minutes at 37C. After removing the medium containing unattached mESCs suspension, mESCs were seeded on 60-mm bacterial-grade Petri dishes at 3×104 cells/cm2 in the same DMEM medium, but in the absence of LIF. EBs formed within 24 hours, and medium was changed every 2 days. Supplementation with 2 µg/ml doxycycline (diluted in sterile PBS and filtered through a 0,2 µm filter unit) (Sigma-Aldrich) and/or 1 µM retinoic acid (RA) (diluted in 0,01% DMSO) (Sigma-Aldrich) were initiated at day 4 and maintained until the required time point for analysis (48h, 4 days or day 8 after doxycycline treatment).

MEFs were derived from wild-type E14,5 mouse embryos following isolation and growth procedures described in (Garfield 2010). Mitotically inactivated MEFs were obtained using 50Gy of gamma irradiation.

### Real-time quantitative PCR

Primers were designed using NetPrimer program and PCR products were confirm by proper melting curves and agarose gel electrophoresis. Values for each gene were normalized to the expression values of *Sdha* and expressed as mean±s.e.m. (of at least three replicates) relative to control untreated samples (without Dox). The primer pairs sequences (forward and reverse) used: Myo7a (5'-TGGGGAGTACAGGTGTGAGA-3'; 5'-CCACAAAGTACTGCTGAGAAGC-3'), Myo6 (5'-GAGAGGCGGATGAAACTTGAGA-3'; 5'-CTTCGGAGTGCCATGTCACC-3'), Otof (5'-CCCAGATCACGGACAGGA-3'; 5'-GCCACCAGCTCTTGATATAGATG-3'), Lhx3 (5'-GCAGTTCCAAGTCCGACAA-3'; 5'-TAGCAGGCCCCATGTCAG-3'), Espin (5'-AGCAGAAGATGCAGGAGGAA-3'; 5'-TTCCGAAGAATGTCTCGTCTC-3'), Cdh23 (5'-AACAGCACAGGCGTGGTGA-3'; 5'-TGGCTGTGACTTGAAGGACTG-3'), Tubb3 (5'-AAGGTAGCCGTGTGTGACATC-3'; 5'-ACCAGGTCATTCATGTTGCTC-3'), Dcx (5'-GACTCAGGTAACGACCAAGACG-3'; 5'-TTCCAGGGCTTGTGGGTGTAG-3'), Barhl2 (5'-GAGAGAAGACCCCCAGAGCG-3'; 5'-TCCCGGTCTCCTTCCTCCTT-3'), Venus (5'-TGAGCAAGGGCGAGGAGC-3'; 5'-AGTCGTGCTGCTTCATGTGGTC-3'), Sdha (5'-CAGTTCCACCCCACAGGTA-3'; 5'-TCTCCACGACACCCTTCTGT-3').

### RNA-seq: data analyses

Sequencing quality of the raw data was checked using FastQC (Andrews 2014) and MultiQC (Ewels et al. 2016). Contaminant adapters were trimmed using cutadapt (Martin 2011). The trimmed reads were pseudo-aligned to cDNA sequences from UCSC's knownGene transcriptome (mm9 assembly) using Kallisto with default settings (Bray et al. 2016). For differential expression analyses, transcript abundances were imported into R and summarised to the gene-level using the tximport workflow (Soneson et al. 2015). Significantly differentially expressed genes (FDR < 0.05 and log2 fold change > 1.5) were identified using the standard DESeq2 package in Bioconductor (Love et al. 2014).The clustering patterns of genes were assessed based on a matrix of the mean of biological replicate samples. The matrix was clustered by use of the pam() function in R. Heatmaps visualizations containing mean normalized counts of each cross-classified gene group were generated using Complex Heatmaps package in Bioconductor (Gu et al. 2016). Lists of genes were analysed for Gene Ontology (GO) term enrichment using topGO package in Bioconductor (Alexa and Rahnenfuhrer 2019).

Gene set enrichment analysis: In vivo hair cell transcriptomes obtained from mouse vestibular and cochlea tissue at embryonic (E16) and postnatal stages (P0, P4 and P7) were downloaded from GEO database, accession number GSE60019. Raw data (FASTQ files) was processed using the same RNA-seq analyses pipeline described above to generate a normalised matrix counts which was imported to GSEA software (Mootha et al. 2003; Subramanian et al. 2005). Defined gene set determined by the above described clustering analyses were also uploaded into the GSEA software to identify statistically significant enrichments between hair cells and non-sensory cell types of the inner ear.

### Luciferase reporter assay

Plasmids used: pCMV5-myc, pGL4.23-6*AtEAM, pCMV-myc-Atoh1, pGL3-6*R21, pCMV5-Gfi1 and pEx-Mm02706-M12. For P19 cells, Thermo Scientific Lipofectamine 3000 Reagent Kit was used for the transfection of 0.5 x $10^5$ cells with 550ng total DNA in 24 well plates. The luciferase assay was conducted 48 hours after the transfection (Promega Dual-Luciferase Reporter Assay System Kit).

The ratio of firefly and Renilla fluorescence intensity was analyzed by GraphPad Prism, calculating the mean value and standard deviation from the three technical repeats in the same group. The mean value of 0 protein group was chosen as the base level, and the fold change of the mean value and deviation from each group was calculated and plotted as a bar chart. To test the significance, the original data from each group went through one-way ANOVA followed by Bonferroni correction, together with Tukey's range test as a parallel check, taking 95% confidence interval.

### GST Pulldown Assay

The GST Pulldown assay was carried out essentially as described in (Nguyen and Goodrich 2006) except lysis and wash buffers contained 5mM β-mercaptoethanol rather than 1mM DTT, and also contained 50μM $ZnSO_4$. Briefly, *E.coli* BL21 plysS cells were transformed with GST or GST-mGfi expression construct (pGEX-KG-GST or pGEX-KG-mGfi1 respectively) and grown overnight in L broth at 25 °C. Protein expression was induced using 1mM IPTG for 5 hours at 25 °C in the presence of 100μM $ZnSO_4$ before harvesting of the GST, or GST-Gfi1 containing bacterial pellet.

The GST alone or GST-Gfi1 was bound to glutathione sepharose and treated with micrococcal nuclease to remove excess nucleic acid. Next a micrococcal nuclease-treated extract containing myc tagged Atoh1 produced from pGBKT7-myc-Atoh1 construct using the TNT Quick Coupled Transcription/Translation system (Promega) was incubated with the GST or GST-Gfi1 complexed to the GSH-sepharose beads for 2 hours at 4 °C. This was followed by extensive washing according to (Nguyen and Goodrich 2006), then the beads were boiled in SDS PAGE sample buffer and the resulting samples run on an SDS PAGE gel, subjected to Western blotting and the blot was probed with anti-myc antibody. A negative control pulldown was carried out in parallel with the unrelated ciliogenesis protein LRRC6 (results not shown).

### Chromatin Immunoprecipitation (ChIP)

6 days old EBs treated with 2 µg/ml doxycycline at day 4 were dissociated using 0.25% trypsin-EDTA (Invitrogen) in PBS and around 15x106 to 20x106 of cells were aliquoted into 15ml falcon tubes for fixation. Cells in each aliquot were fixed at room temperature for 12 minutes on 1% of formaldehyde solution (37% formaldehyde, Sigma-Aldrich). Formaldehyde was quenched with glycine (final 0.125 M) for 5 minutes at room temperature. Samples where then washed once with cold PBS containing protease inhibitors (cOmplete protease inhibitor cocktail, Roche). After, cells were centrifuged at 500 g and pellets were snap frozen on dry ice followed by storage at -80C until further use. The pellet for each sample (~20x106 of cells) was thawed and nuclei were isolated by re-suspension in lysis buffer (5 mM Pipes pH 8.0, 85 mM KCl, 1% NP40) with fresh protease inhibitors (cOmplete protease inhibitor cocktail, Roche) for 20 minutes on ice followed by brief vortexing and centrifugation at 500 g for 10 minutes at 4C. The nuclear pellet was re-suspended in 300 ul IP buffer (0.5% SDS, 1% Triton, 2 mM EDTA, 20 mM Tris-HCl pH 8.0, 150 mM NaCl, fresh protease inhibitors) and sheared by sonication (BioRuptor Pico, Diagenode) down to ~200-300 bp fragments using a total of 5-14 cycles of: 10 pulses of sonication on the ‘‘high’’ setting, each followed by 30 s off. Samples were then diluted in IP buffer without SDS, then 10 mg of the following antibodies was added for overnight incubation at 4C on a rotator: Monoclonal anti-flag M2 antibody (F1804, Sigma); Myc tag antibody-ChIP Grade (ab9132, Abcam); Anti-Gfi1 antibody (ab21061, Abcam). Dynabeads-ProteinG (Life Technologies) were blocked for 1-hour in 0.5% BSA in IP buffer with protease inhibitors, prior to incubation with the sonicated antibody bound chromatin suspensions. Bead-chromatin complexes were then serially washed for 5 minutes each with the following solutions: IP buffer (150 mM NaCl), followed by IP buffer with high salt concentration (500 mM NaCl), then 1 wash with washing-buffer (10 mM Tris-HCl pH 8.0, 0.25 M LiCl, 0.5% NP40, 0.5% Na-Deoxycholate, 1 mM EDTA), followed by 2 washes with TE (10 mM Tris, 1 mM EDTA). The DNA-protein complex was then eluted from beads by incubation with 100 ml of elution buffer (1% SDS, 10 mM EDTA, 50 mM Tris-HCl pH 8.0) for 30 minutes at 65C, vortexing every 10 minutes, followed by magnet extraction of beads. The beads were re-washed with 150 ml TE with 1% SDS, magnet extracted, and the TE with remaining DNA solution was added to the eluted samples, followed by 65C overnight incubation to reverse crosslink the DNA-protein complexes. The dissociated DNA and protein solution was then treated with 4U of proteinase K (NEB) at 37C for 2 hours. DNA was isolated with SeraMag beads, using 450 ml SeraMag bead solution and 225 ml of 30% PEG in 1.25 M NaCl (Rohland and Reich 2012). After bead purification, DNA was re-suspended in 30 µl of ddH2O (DNase/RNase free) and for each ChIP sample, DNA concentration was estimated using Qubit fluorometric quantification (Invitrogen) before being stored at 80C.

### ChIP-seq: library preparation

8-10 ng of ChIP DNA was used for each sample. DNA ends were blunted by treatment with 5 µl T4 DNA ligase buffer (NEB), 2 µL 10 mM dNTP’s, 0.5 µl end repair mix (0.72 U T4 DNA polymerase, 0.24 U Klenow Fragment, 2.4 U T4 DNA Polynucleotide Kinase), up to 50 µl with ddH2O (DNase/RNase free). Samples were incubated for 30 minutes at 20C, and then purified with addition of 50ul

SeraMag bead solution and 50 µl 30% PEG solution (in 1.25 M NaCl). DNA was eluted in 16.5 ul of TE/10 (10 mM TrisCl pH 8.0, 0.1 mM EDTA). DNA was then A-tailed at the 3' ends by treating the eluate with 2 µl 10X NEB Buffer 2, 1 µl 4 mM dATP, 0.5 µl Klenow 3' to 5' exonuclease minus (NEB) in a total volume of 20 µl. Reaction was incubated at 37C for 30 minutes. Before adapter ligation, TruSeq adapters (Illumina) were first annealed by re-suspending each adapter at 100 mM (in 10 mM Tris-HCl pH7.8, 0.1 mM EDTA pH 8.0, 50 mM NaCl) and mixing them at a 1:1 ratio. Adaptor annealing occurred using a program of 2 min at 95C, 70 cycles of 30 s (95C decreasing by 1C each cycle). Annealed adaptors were diluted 1:200 (0.25 mM final) and then 1 µl was used for ligation reaction containing 20 µl of the A-tailed DNA, 1.5 µl Quick Ligase (2,000 U/ml NEB) and 2.5 µl of H2O. The reaction was incubated 20 minutes at room temperature and stopped by addition of 5 µL of 0.5 M EDTA pH 8.0. Next, DNA was purified using 50 µl SeraMag bead mix and 50 µl 30% PEG (in 1.25 M NaCl) and eluted in 15.5 µl of TE/10. Next, the adapter ligated DNA was amplified using the following protocol: 15 µl adaptor-ligated DNA, 1 µl TruSeq primer cocktail (0.25 mM), 15 µl 2X Kapa HiFi HotStart ready mix. The Libraries were amplified with the following protocol: 45 s at 98C, 5 cycles of (15 s at 98C, 30 s at 63C, 30 s at 72C), 1 minute of 72C, hold at 4C. After pre-amplification of the library, a size selection step was performed to obtain DNA fragments between 300 and 500 bp using the following procedure: 0.9X beads were added to ChIP DNA After 15 minutes incubation and 10 minutes magnet extraction, the supernatant was transferred to new tube and beads discarded. Then 0.2X beads were added to the solution, incubated 15 minutes followed by 10 minutes magnet extraction and beads re-suspension in 11.5 µl TE/10. 11 µl of pre-amplified and size selected DNA was further amplified with 1 µl TruSeq primer cocktail, 20 µl 2X Kapa HiFi HotStart ready mix, and 8 µl H2O using a PCR program of 45 s at 98C, 9-11 cycles of (15 s 98C, 30 s 63C, 30 s 72C), 1-minute 72C. After amplification, the DNA was isolated using SeraMag bead purification. Libraries were assessed using Agilent High Sensitivity D1000 ScreenTape System on an Agilent 2200 TapeStation. Finally, libraries were pooled at equal concentration and sequenced with Edinburgh Genomics on the Illumina HiSeq 4000 platform.

### ChIP-seq: data analyses

Reads quality was checked using using FastQC (Andrews 2014) and MultiQC (Ewels et al. 2016). Illumina contaminant adapter sequences were removed using Trimmomatic (Bolger et al. 2014). Reads were aligned to the UCSC mm9 assembly of the mouse genome using Bowtie 2 with the very-sensitive option (Langmead and Salzberg 2012). For downstream analysis, PCR duplicates reads were removed using Picard Tool's MarkDuplicates command (Broad Institute) and were filtered for MAPQ > = 40 using SAMtools (Li et al. 2009). Reads aligned to ENCODE blacklist regions regions (Consortium 2012) were removed using BEDTools (Quinlan and Hall 2010). Unmapped, secondary, and supplemental read alignments were filtered using SAMtools. Peak calling was performed by first estimating the fragment size using cross-correlation analysis from the phantompeakqualtools software package (Landt et al. 2012). Peaks were called using MACS 2 on merged sample replicates against merged control replicates (Zhang et al. 2008). The following arguments were used: --g mm --keep-dup all -s <read length> --nomodel --shift 0 --extsize <fragment size> -p 0.05 -B --SPMR.

<u>Read coverage visualisation</u>: DeepTools suite (Ramírez et al. 2016) was employed for read coverage visualisation. Normalized input-subtracted read coverage was calculated using merged control and samples with the ratio subtract and normalize to RPKM options. Heatmaps were generated using plotHeatmap command with the normalized input-subtracted read coverage and peaks called from

the merged sample replicates. The DiffBind software package from the Bioconductor project (Ross-Innes et al. 2012) was used for correlation and principal component analyses of all ChIP-seq data.

De novo motif analysis: was performed with the findMotifsGenome tool of the HOMER (Heinz et al. 2010) suite searching for 250 bp around the peak summit. The background was generated by the HOMER software and only motifs with P-value < $1x10^{-50}$ were consider significant. The tool annotatePeaks.pl was also used to find specific motifs 250 bp around the peak summit.

Overlapping peaks and peak to gene assignment: Peaks were considered to be overlapping when there was at least 1 bp of overlap between a 300bp region centered on each peak summit. These overlapped peaks were obtained using mergePeaks tools from the HOMER software. Overlapping and unique peaks were annotated to the nearest TSS using ChIPseeker (Yu et al. 2015) package in Bioconductor. For most of the analyses, only peaks associated with differentially expressed genes were used for functional analyses that were carry out with clusterProfiler package (Yu et al. 2012) and topGO in Bioconductor (Alexa and Rahnenfuhrer 2019). Venn diagrams and histograms were generated using the package Vennerable (Swinton 2009) and ggplot2 in the R statistical environment.
